# Deviation from Power-Law Distribution when Scaling the Distribution of Marine Plankton Folds from Genomes to Communities

**DOI:** 10.1101/2025.03.25.645231

**Authors:** Lucas Pavlovic, Caroline Vernette, Emanuele Pigani, Louis Joigneaux, Alexandre Labesse, Janaina Rigonato, Daniele Iudicone, Magali Lescot, Youri Timsit, Olivier Jaillon

## Abstract

At different scales of living systems, biological entities appear to follow scaling laws, such as power laws, which are often explained from stochastic mechanisms. This is the case for the number of species in a community and the number of genes or protein folds in a genome. Resulting from evolutionary processes combining gene family duplications and expansions with selective pressures, the distribution of protein folds systematically follows a power law in all individually observed genomes. A small number of folds are highly prevalent, while the majority of folds appear only once per genome. However, previous studies on fold occurrence have focused on individual genomes, isolated from their community contexts. In the oceans, plankton communities consist of complex assemblages of species, each exhibiting variable relative abundances. We investigated the consequences of this variability on the composition and distribution of folds by considering the relative abundance of species. By annotating folds to genes of environmental genomes of plankton collected by the Tara Oceans expedition, we show that the relative abundance of folds deviates from the classical power law and instead follows a Type II Pareto distribution. This model, typically observed in other complex organizations such as economics, allows us to classify different categories of folds that exhibit biogeographical differences. Our results show that scaling fold distributions from individual genomes to species communities lead to a deviation from the expected behavior of simple power-law relationship towards a more complex model. This phenomenon could be linked with the variable complexity of marine planktonic ecosystems.

## Introduction

Scaling laws describe the distribution of biological entity abundances across various biological systems, from genomes to ecosystems. For example, in marine ecosystems, the distribution of species abundances typically follows heavy-tailed distributions (log-normal and power laws for instance), with a small number of abundant, dominant species and a large number of rare, non-dominant ones [1], [2], [3], [4]. At a much smaller scale, the distribution of protein folds in genomes also follows a power law, with a few highly abundant *superfolds* (SFs) and many rare folds, called *unifolds* [5], [6], [7], [8]. Bridging scales is a long-standing challenge, aiming to decipher cross-scale interactions to better understand and model complex living systems. In this study, we address this question by analyzing the distribution and abundance of protein folds in marine planktonic ecosystems. For this purpose, we have attempted to bridge knowledge from structural biology and ecology by analyzing the fold abundances of metagenomic data collected during the *Tara* Oceans (TO) expedition [9].

Plankton is one of the most diverse microbiome on the planet, and has the particularity of being passively carried by currents [10]. Along their journey, planktonic species face important changes of environmental parameters (day length, water temperature, nutrient concentration, …) in period of times that range from days to years [11]. Depending on their body size, their distribution in the oceans is biogeographically structured either by latitude [12], [13], biomes, physics or Longhurst biogeochemical provinces [11], [14], [15]. Some species are also cosmopolitan and abundant in every sampled basin [16]. The plankton proteome, the collection of proteins that are expressed in the plankton cells, is a key system responsible for life behaviour at the scale of individuals, but also at the scale of community considering complex biotic interactions [17], [18].

At the nanometer scale, protein domains adopt a fold which is the three-dimensional pathway of the polypeptide chain skeleton. The diversity of globular folds is considered limited, between 1350 and 4000 different folds [8], [19], [20]. There are currently 1472 different folds in CATH [21], [22], 1257 in SCOPe [23], [24] and 1562 in SCOP2 [25], [26]. Although the number of protein folds catalogued in databases has remained relatively stable for many years (despite the exponential growth in genomic and proteomic data), AlphaFold models have increased the CATH database with nearly 200 new folds [27]. The Encyclopedia of Domains, based on the AlphaFold Protein Structure Database, might also reveal thousands of putative new folds [28]. Folds usages vary across the tree of life and genomes [5], [29], [30], [31], [32]. While some folds are universal, others are lineage-specific and can be considered as synapomorphies [33]. The emergence of novel folds was particularly important during the evolution of Eukaryotes since 10-20% of the fold diversity is specific to that domain of life [33], [34]. Moreover, it contains the highest number of multidomain proteins with unique fold combinations, which usually represent at least two-third of a proteome and up to 80% in Metazoans [35], [36]. The functional versatility of each fold is variable [37]. The SFs, like the Rossmann fold, are highly occurring in genomes partly because they display an extremely diversity of functions [35], [38], [39], [40]. Previous studies have shown that fold occurrences follow a power-law distribution in proteomes of Eukaryotes, Prokaryotes and Viruses [5], [6], [41], [42], which results from the fact that a little number of SFs gathers most of the fold occurrences while a high diversity of *unifolds* are adopted only once or twice.

While these studies clearly demonstrated that fold usage in genomes results from long-term evolutionary processes, the question addressed here is: “is the distribution of protein folds at the level of plankton communities similar to that of genomes, or does it differ according to community or environments?”. Indeed, the different categories of protein domains exhibit distinct biophysical and functional properties. For instance, depending on their content in α-helices or β-sheets, proteins differ in folding dynamics, stability, allosteric properties and even robustness to mutations [43], [44], [45], [46], [47], [48], [49]. These variations strongly suggest that biophysical constraints may play a key role in shaping the distribution of folds across different environments. It is known for example that proteomes from species with different ecological preferences (mainly thermophile or psychrophile) have different properties [50]. It has been recently shown that persistent environmental changes may also induces changes of the fold usage in fungal proteomes [51]. The impact of the environment on the proteome was also observed in the comparison of mesophilic and psychrophilic diatom species. The study revealed that the elevated zinc concentrations that characterize the Antarctic region resulted in a greater abundance of zinc-binding proteins in Antarctic diatoms [52]. To our knowledge, fold usage has never been investigated at the level of sampled marine planktonic communities, composed of barely known organisms and at a wide geographical scale.

In this study, we investigate the fold diversity and usage of planktonic communities with two main objectives: first, to determine the variability of fold repertoires in lineages distant from model organisms; second, to examine the compositional and biogeographic properties of fold repertoire assemblies in natural communities. To do so, the proteomes of eukaryotic, prokaryotic and *Nucleocytoviricota* (NCLDV) MAGs from the TO expedition [53] were structurally annotated using CATH in order to characterize the distribution of the folds in environmental genomes and communities. We show that at the individual genome scale, fold usage follows a power-law distribution as expected. Besides, by leveraging the occurrence of folds in proteomes by the relative abundances of genomes within their communities, we reveal the distribution of folds in a community sample composed of assemblages of multiple species with varying abundances. While the most common folds still follow a power-law distribution, the less common and intermediate folds gradually deviate from it, resulting in an overall distribution that transitions toward a Pareto-like pattern. This indicates that different principles govern fold occurrences within a genome and fold abundance within a community. These findings provide new insights into how ecological constraints shape the evolution of proteomes.

## Results

### 1) Fold occurrences in environmental genomes of plankton

#### 1.1 Fold occurrences in genomes follow a power law

The quantitative distributions of fold occurrence values (OVs) in the environmental plankton genomes were studied. We used a previously published collection of approximately 700 eukaryotes, 1,900 prokaryotes and 31,000 NCLDV [53],[54]. Due to the incomplete reconstruction of these genomes, with an average completeness of 30%, we first controlled their suitability for our study. To this end, we compared the distribution of fold occurrences between eukaryotic MAGs and the proteomes of reference (RPs) from the EukProt database [55]. The distributions obtained were globally similar (Supplementary Fig. 1). The main differences suggest that genome incompleteness primarily affects the detection of rare folds and, importantly our study, does not alter the shape of the distribution. This confirms that the incompleteness of Eukaryote MAGs is unlikely to bias our main conclusions.

The distribution of fold OVs in our datasets systematically have a power-law behavior in Eukaryotes, Bacteria, Archaea as well as in NCLDVs, in a manner consistent with previous studies [5], [6], [42] (Supplementary Fig. 2; Supplementary Fig. 3). It confirms that in environmental genomes, a small number of folds (the SFs) have such high OVs that they account for the majority of the total fold occurrences. For example, in the eukaryotic MAGs, the Rossmann fold (3.40.50) alone represents around 15% of the total occurrences, which is roughly the proportion identified in other studies [40]. Conversely, a large number of different folds are present only a few times.

#### 1.2 Power-law parameters vary among lineages

It is interesting to note significant differences in the shape and parameters of the laws modeling the fold occurrence distributions across domains of life but also at a lower taxonomic rank in Eukaryotes and Bacteria (Supplementary Fig. 2-4). Eukaryotes exhibit a higher proportion of frequent folds compared with prokaryotes, likely due to their larger genomes and higher gene duplication rates [5], [6]. Bacteria and Archaea share more similar fold distributions, suggesting common evolutionary constraints. Among eukaryotic phyla, the power law modelling the distribution of fold OVs in Arthropoda and Choanoflagellata have parameters that deviate compared with the ones of unikonts. Accordingly, power-law parameters could serve as indicators for major phyla, at least in eukaryotes. These findings confirm, on environmental genomes that encompass a vast part of the diversity of eukaryotes, how genome evolution, in particular duplication, shapes proteome diversity across taxa. The contrast between eukaryotes and prokaryotes suggests that genome complexity drives structural diversity, while the similarities between Bacteria and Archaea point to conserved constraints in fold evolution.

### 2) Mapping fold abundances in global ocean

The relative abundances values (AVs) of folds within marine planktonic communities were calculated by taking into account the relative abundance of their associated genomes in metagenomes. Thus, the calculated fold abundances depend on both fold occurrence in genome and the abundances of the MAGs in the ecosystem (Mat. and Meth.) The global representation of the fold compositions across sampling sites visually reveals several categories of folds based on their abundances (Fig. 1A). For comparison purposes, the same visualisation but with MAG AVs is available in Supplementary Fig. 5B. While some folds are highly abundant everywhere (folds on the left of Fig. 1A), some have variable AVs and are present almost everywhere, in some cases with punctual absence in one or two stations that are probably artifactual; others are rare and absent in a high number of station. While some highly abundant folds correspond to well-known SFs like the Rossmann one, others are not, such as the alpha-horseshoe (1.25.40). The fact that only a very small fraction of the fold diversity (roughly 8%) holds by far the highest abundance values indicates that the distribution of AVs is probably power-law like, as observed for the OVs above. The similarities of TO stations in terms of fold composition do not appear to be geographically structured (Fig. 1B).

**Figure 1.**
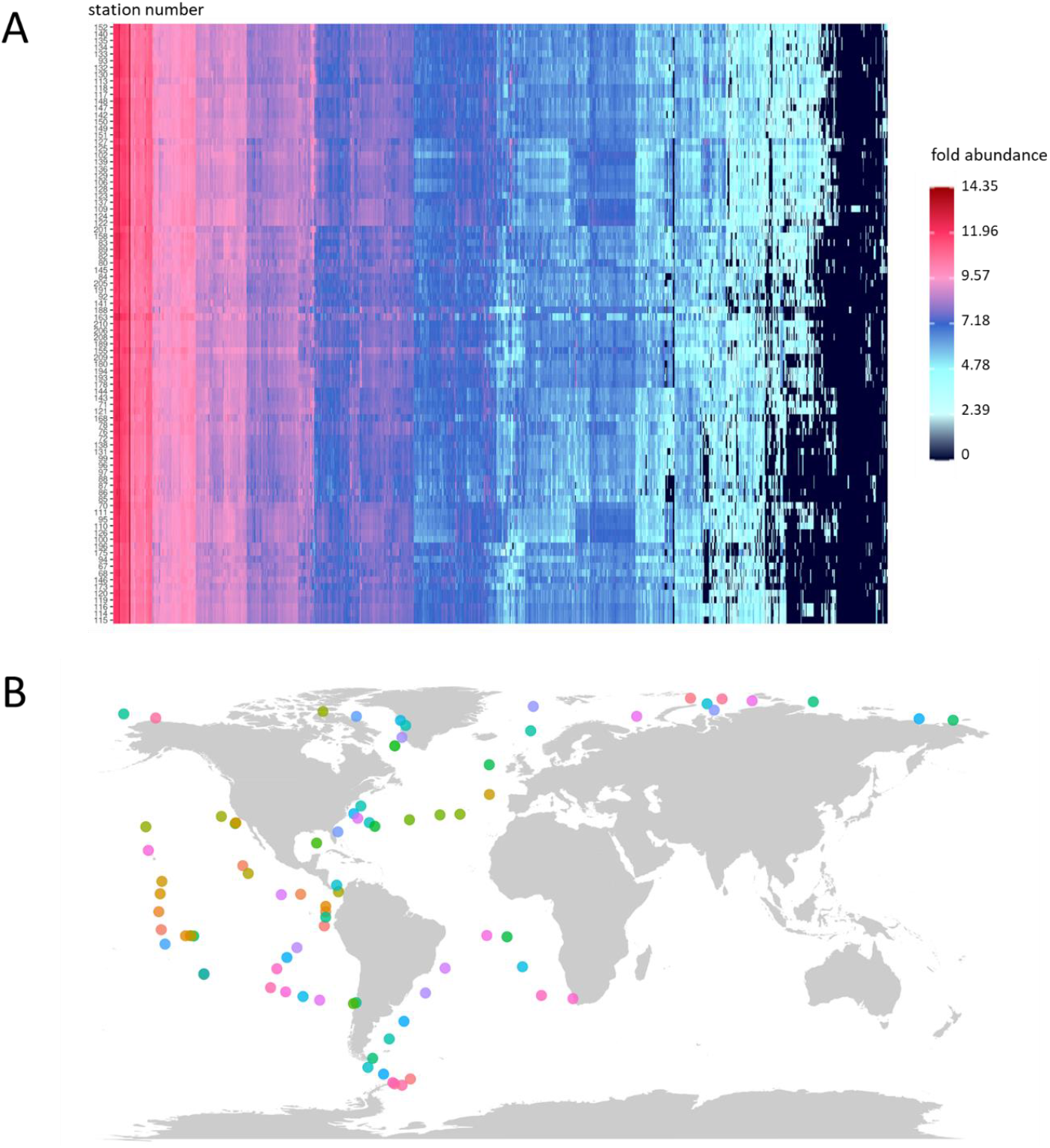
Distribution of the 908 folds in the surface and 0.8-2000µm size fraction of 89 TO stations. **(A)** Heatmap of the Centered Log-Ratio (CLR) translated AVs of the protein folds in the TO stations. Station numbers are indicated on the y-axis. They are ordered following their clustering based on the AVs of the folds. The 908 folds are on the x-axis (not displayed). They are ordered based on the reciprocal clustering. Large clusters were then manually reordered. A version of this figure with supplementary layers both on the y- and x-axis can be found in Supplementary Fig. 5A. **(B)** Map of the sampling sites with colours reflecting their degree of similarities. A PCoA was calculated based on the same distance matrix as in **(A)**. The coordinates of the stations in the PCoA space were then translated into a Red-Green-Blue (RGB) code, which was used to colour TO stations on a world map. Sites with similar colours are close in the PCoA space and accordingly have similarity in terms of fold composition.

### 3) Pareto law describes fold abundance distribution in marine ecosystems

#### 3.1 Fold abundance distributions deviate from power law and follow a Pareto law

A comparison between OV and AV distributions of folds in the TO stations highlights a significant impact of the MAG AVs on the fold AVs in the environment. Notably, the distribution of fold AVs sometimes deviate significantly from the power-law model, apparently because of the low AVs, resulting in a lower regression coefficient. In these cases, the overall abundance distribution seems to follow two distinct regimes separated around a transition point (Fig. 2A).

**Figure 2.**
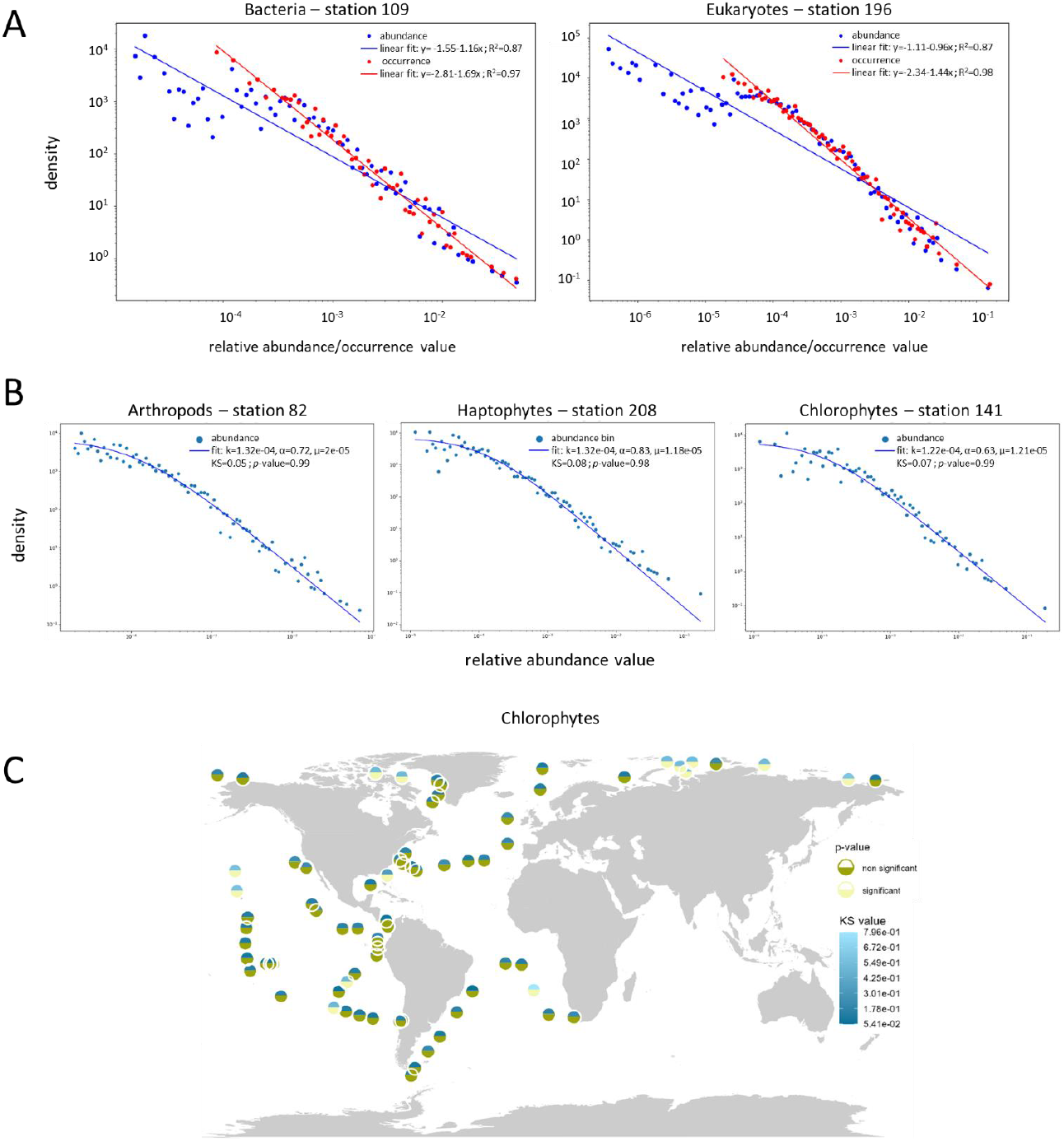
Pareto type II distribution of the folds in the environment. **(A)** Example of linear fits for the fold OVs (in red) and AVs (in blue) in Bacteria (left) and Eukaryotes (right) in sampling stations 109 and 196, respectively. The regression equations and associated R^2^ are displayed on the top right corner of each fit. **(B)** Pareto type II (PII) fits on the fold abundance of three selected eukaryotic phyla (left: Arthropoda; middle: Haptophyta; right: Chlorophyta) in three selected sampling stations (left: 82; middle: 208; right: 141). Values of the parameter of the equation (k, α, µ), of the Kolmogorov-Smirnov (KS) test and associated *p*-value are on the top right corner of each fit. **(C)** Map of the result of the KS tests for the distribution of fold AVs of Chlorophyta. Each dot is a TO station. The upper colour (shades of blue) corresponds to the value of the KS statistic (the lower, the darker, the better the fit). The lower colour (shades of green) indicates whether the test is significant or not, under the *H*^*0*^ *“the observed data follow the Pareto type II model of given parameters”*. Non-significant tests (*p*-value>0.01) are indicated in dark green, significant ones in light green (a significant test rejects the hypothesis of a PII fits whereas a non-significant indicates a compatibility with a PII).

To better account for the deviation of AV distributions from the power-law model, we tested alternative models within the Pareto family, which are generally well-suited for heavy-tailed data distributions. Among these models, the Pareto Type II (PII) law provided the best balance between accuracy and complexity. In the PII distribution, two regimes are distinguished (left-hand and right-hand), which do not have the same slope in log-log representation. In the complete equation, the scale parameter *k* represents the value on the x-axis that separates the two regimes. We used this law to model the AV distributions and validated the models using Kolmogorov-Smirnov (KS) tests and their associated *p*-values (Mat. and Meth.). Considering all eukaryotic MAGs or, independently, all bacterial MAGs, the models obtained were satisfactory for almost all samples (Supplementary Fig. 6). These results indicate, first, that the deviation from the power-law model is unlikely to result only from a bias specific to eukaryotes; and second, that the transition from a power law to a PII model occurs when considering the distribution of folds at the scale of a community, which accounts for the relative abundances of different species and, to a certain extent, their ecological dynamics, rather than at the scale of individual genomes.

We mentioned above that, at the genome scale, certain power-law parameters of the fold distribution appear to serve as indicators for major phyla. When considering the relative AVs of genomes within communities, the distributions of KS values and p-values indicate that the properties of PII laws vary across stations and eukaryotic phyla (Fig. 2B, Supplementary Fig. 8A)

#### 3.2 variability of the KS values among lineages and environmental conditions

Given their variability across phyla and stations, the parameters of the PII law may thus reflect the ecological characteristics of the sampled environments. Since MAG AVs at a near-community scale would be dependent on local ecological processes, fold AVs are influenced not only by these processes but also by the genomic properties of the MAGs within the community. PII fits associated with a *p*-value superior to 0.01 and KS values inferior to 0.2 were considered “good” in the rest of the analysis (that is, the PII fits cannot be statistically rejected by the KS test).

PII parameters and results of the KS tests vary according to phyla and stations (Fig. 2B, Supplementary Fig. 8A), especially in terms of proportion of stations where the distribution of fold AVs follows a PII. These proportion are calculated as a ratio of the number of stations with a “good” model on the total number of stations where MAGs of the phylum are present. Among eukaryotes, Chlorophyta, Arthropoda and Haptophyta are the phyla with the highest proportion of “good” PII fits, with 90, 85 and 82%, respectively. These proportions were lower in Bacillariophyta and MAST-4 (51 and 44% respectively), and even more in Choanoflagellata (only 10%).

To better understand this phenomenon, the results of the KS tests were displayed on a world map (Fig. 2C; Supplementary Fig. 7). When considering all eukaryote MAGs together, the distribution of fold AVs does not follow the PII law in only two stations: 196 and 68. The comparison of the geographical repartition of “good” fits at the scale of eukaryotic phyla provides a first possible indication of an environmental influence on fold distribution. Indeed, “good” fits in Chlorophyta and Haptophyta are mostly observed outside of the poles contrary to Bacillariophyta, probably resulting in part from a distinct fold repertoire between polar and non-polar species [14], [15]. Interestingly, the “good” fits in Bacillariophyta are linked with the complexity of the community. Indeed, they tend to be observed in stations with a high α-diversity and richness, which is surprisingly not the case with Chlorophyta and Haptophyta (Supplementary Fig. 8B-C)

### 4) Abundance fold categories

In the PII law, the *k* parameter allows for an objective distinction between different regions of the fold distribution, as indicated before. We used it to classify folds into three abundance categories (Supplementary Fig. 9). Folds in the first category have AVs consistently below *k* in every sampled station, whereas folds in the second category always have AVs above *k* across all sampled stations. We defined an intermediate category for folds which AVs can be either below or above *k* depending on the station. The core of each of these categories corresponds to the folds that belong to the same category in the six phyla.

Depending on the phylum, the highest number of folds is found either in the first category (Bacillariophyta, Choanoflagellata, MAST-4) or the intermediate one (Chlorophyta, Haptophyta, Athropoda). The size of the second category is relatively stable across phyla (Supplementary Fig. 10). In the different categories, around 30% of folds are Mostly Alpha, 20% of folds are Mostly Beta, 35% Alpha Beta and between 0 and 10% Few Secondary Structures and Specials (there are no Special folds in the 2^nd^ category (Supplementary Fig. 11). These proportions are close from the ones observed at the scale of proteomes (Supplementary Fig. 3), indicating that the abundance categories are not particularly enriched in any Class or Architecture and their fold composition is representative of the fold composition of eukaryotes in general.

#### 4.2 Environmental parameters that affect the fold distributions

To better understand the ecological significance of the abundance categories of folds, the AVs of the folds in each category and phylum was displayed (Fig. 3A). It revealed that the folds of the 1^st^ category have the lowest abundances, followed by the ones of the intermediate category and eventually the ones of the 2^nd^ category. Accordingly, we rename the 1^st^ and 2^nd^ category “rare” and “abundant” for the rest of the analysis. The order of the folds in Fig. 1A is also globally consistent with these names (Supplementary Fig. 13)

**Figure 3.**
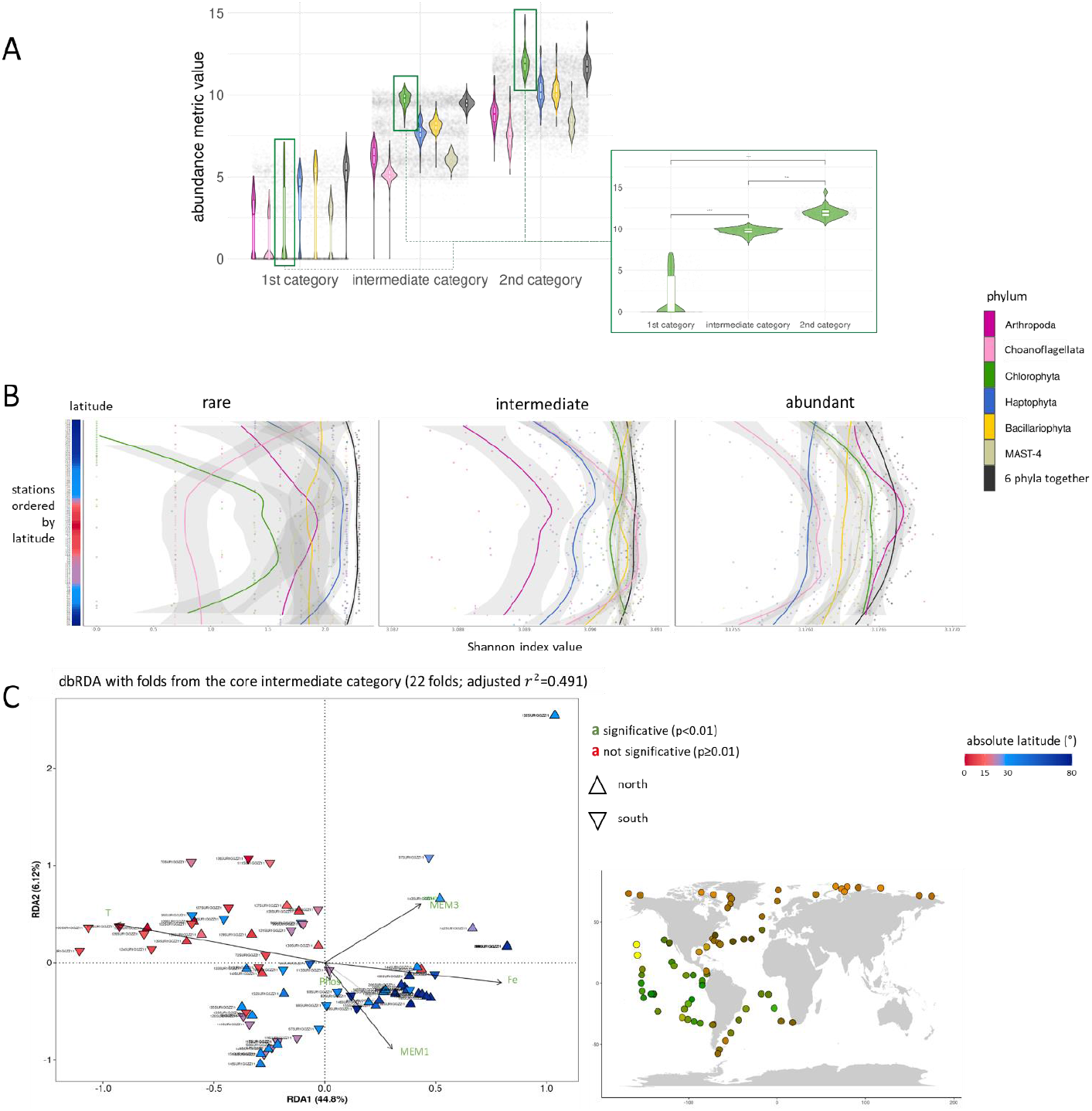
Biogeographic properties of the distribution of the eukaryotic MAGs folds in the global ocean. **(A)** AVs of the folds of each category and phylum. The violin plots are organized by category (from left to right: first, intermediate and second) and within category, by phylum (from left to right: Arthropoda, Choanoflagellata, Chlorophyta, Haptophyta, Bacillariophyta, MAST-4 and the six phyla together). Statistical significance within and between groups was tested with a Wilcoxon test, as exemplified on the Chlorophyta case (the results on other phyla are available in Supplementary Fig. 12) (***: *p*-value < 0.01). **(B)** Shannon index of the folds from the core of the different abundance categories (from left to right: rare, intermediate, abundant) per phylum. The stations are indicated on the y-axis, ordered by latitude. The values of the Shannon index are indicated on x-axis. Each colour represents a phylum. The grey zone surrounding the lines indicate the standard error. **(C)** Distance-based ReDundancy Analysis (dbRDA) biplot (on the left) and corresponding Red-Green (RG) map (on the right) of the folds from the core of the intermediate category in Chlorophyta. The subtitle indicates the number of folds in brackets, followed by the value of the adjusted coefficient of determination *r*^2^ (proportion of variance of the fold matrix explained by the selected variables). In the biplots, the coloured triangles represent the stations. Their colour correspond to the absolute latitude. Triangles pointing upwards correspond to stations above the equator. Downward-pointing triangles correspond to stations below the equator. Black arrows correspond to statistically significant explanatory variables, and point toward names in green. They are transparent otherwise and point toward names in red. MEM: Moran Eigenvector Map. T_woa: mean annual sea surface temperature (SST) extracted from the WOA [56]. Sd_T_woa: standard deviation of the mean annual SST extracted from the WOA. Phos_woa: Phosphate concentration extracted from the WOA. Fe: Iron concentration from the PISCES-V.2 model [15], [57]. The coordinates of the stations in the two first dimensions of the dbRDA space were converted to a RG code that was used to colour the corresponding stations in a world map. Other results for the Chlorophyta are available in Supplementary Fig. 17-18. Results for the other phyla are summarized in Supplementary Table 3-5.

Next, we verified if those differences of AVs between categories were also linked with differences in α-diversity. Using the Shannon index, we observed an almost systematic decay of α-diversity toward the poles (Fig. 3B; Supplementary Fig. 14-16; Supplementary Table 2), reminiscent of results from previous studies that observed such a pattern at the species level [12]. To confirm the polar effect in the structuration of these repartitions, we performed distance-based ReDundancy Analysis (dbRDA) on each category-phylum combinations. They yielded relatively low determination coefficients, at the exception of intermediate and abundant folds and their core in Chlorophyta (Fig. 3C; Supplementary Fig. 17-18; Supplementary Table 3-4). In these two analysis, median temperature exhibited the strongest association with the observed structuring, followed by the iron concentration and at least one MEM, indicating the importance of the spatial distribution of the sampling stations (Supplementary Table 5). For comparison, the distribution of Chlorophyta MAGs in the TO stations is not biogeographically structured, confirming a stronger biogeographic structuring at the scale of the folds in Chlorophyta than at the level of environmental genomes (Supplementary Fig. 19).

Consequently, in these specific cases, the environmental conditions likely participate to some extent to the observed structuring of the distribution of the folds.

## Discussion

### Variability of the power-law models of fold distribution in individual genomes

CATH can structurally annotate a protein sequence only if a sufficiently close homolog already exists in Uniprot and Ensembl. Many NCLDVS and many eukaryotic taxa lack such representation, leading to variable proportions of unannotated proteomes. However, this non-random variability is unlikely to alter the results given the scale free property of the power-law distribution. In theory, this property implies that a power-law distribution should be preserved in any randomly drawn subset of an ensemble that follows this law, regardless of its size.

The differences observed between RGs and MAGs in eucaryotes are both quantitative and qualitative, and are, to some extent, consistent with the taxonomic composition of the respective database. For instance, the only Metazoans present in the MAG database are Copepods and Tunicates, whereas EukProt includes a broader representation of metazoan diversity. Metazoans, particularly Vertebrates, are known to have experienced intense gene duplication events throughout their evolution, which has, in turn, influenced the OVs of folds in their genomes [42], [58].

On the other hand, single copy genes and rare genes are less prone to undergo duplication. This pattern has been described has being related to the phenomenon of preferential attachment, which results in power-law distribution [6], [8]. The more intense the duplications of large gene families, the lower the slope of the regression curve on a double logarithmic scale, indicating a reduced frequency difference between small and large gene families.

The properties of folds further amplify this trend, as all genes within a given family generally share the same fold, and unrelated gene families can also converge on the same fold. This phenomenon is likely responsible for the differences observed between the regression slopes of Eukaryote MAGs and RPs, with the latter exhibiting a slightly lower slope. This difference aligns with the fact that RGs are enriched in genomes with high rates of gene duplication.

From a qualitative point of view, folds belonging to Class 4 and Class 6 are moderately more frequent in the EukProt structuromes. These classes correspond to non-globular folds with fewer secondary structural elements. In eucaryotes, they are frequently associated with signalling and regulatory pathways [59]. The diversification of these functions has been particularly prominent in metazoans [34], [58]. It is therefore coherent to find these folds enriched in EukProt compared to MAGs.

### Significance of the transition from a power law in individual genomes to a PII law in communities for modeling fold distributions

By attempting to build a bridge between structural biology, environmental genomics and marine ecology, this study provides new perspectives for these disciplines. This work shows that the protein fold distributions in genomes as well as in communities, in particular the mathematical parameters of their models (regression coefficient for the linear models and *k* for the PII model), can constitute an interesting marker for communities of marine plankton. Despite certain methodological limitations, such as the incompleteness of the MAGs, their uneven depiction of marine planktonic communities and the heterogeneity of the CATH annotation rate, this analysis demonstrates that the power-law distribution of fold occurrences, previously observed in individual genomes from model organisms is also present in individual environmental genomes. Moreover, our study significantly expands knowledge on fold occurrences in genomes, extending it to species that are underrepresented or absent from reference genome databases [40], [41], [42].

An important finding is that in contrast to the fold occurrence distribution, the models of distribution of fold abundances deviate significantly from the power law and follow a PII law in the global ocean. What does this model tell us? In the PII law, the two observed slopes—one shallower than the other— indicate the presence of two distinct regimes. For low fold abundances, the decline in the abundance distribution is less steep than for higher values. This phenomenon suggests a form of attenuation or damping of the power law for low-abundance folds. The PII distribution thus reveals that, within each environment, the AVs of genomes constituting a community smooth or moderate the sharp nature of the power-law distribution that typically characterizes individual genomes isolated from their ecological context. This is because a given fold may exhibit varying occurrences across the genomes of different planktonic species that each have a relative abundance that is partially impacted by its biotic and abiotic interactions, as well as by ecological drift. Within a community, this variability can lead to a reordering of fold occurrence ranks, particularly for folds with low-to medium-occurrence. This “shuffling” likely drives the deviation from a power law toward a PII, as it can either amplify or reduce the abundance of certain fold categories. Once again, this analysis underscores that *“the whole is greater than the sum of its parts”*, particularly when transitioning from the genomic to a higher ecosystem level. Interestingly, such deviations from the power law has been observed in other systems [2], [60], [61], [62], [63]. Our analysis can therefore provide the mathematical and conceptual basis for a better understanding of how systems deviate from the power laws.

More globally, these results highlight the necessity of considering biological phenomena holistically. From an ecological and evolutionary point of view, what we observe is likely the result of the combination of two different timescales governed by different evolutionary forces shaping a biological system. The power law reflects long-term evolutionary processes shaping genomes over millions of years, via events of genomic duplications [5], [6] while the PII law, in contrast, captures the modulation of communities over shorter timescales, driven by biotic and abiotic interactions within ecosystems — ranging from hours in response to meteorological or biotic events to years in response to oceanographic physical processes. Therefore, the composition of community-level fold results from two types of processes occurring at vastly different temporal scales. These findings provide insights into the interplay between evolutionary and ecological strategies that organisms use to adapt over long timescales as well as how communities continuously adjust on timescales as short as a day or less. From another perspective, these findings reveal a link between phenotype and genotype at the ecosystem level.

The attenuation of the sharpness of power-law models—where *the poorest become slightly less poor* within an ecosystem—is reminiscent of a recent study on the ubiquitous distribution of non-dominant plankton species in the global ocean [1]. The idea that ecological interactions modulate abundance distributions and smooth out inequalities originates from macroecology and ecological network theory. Studies in community ecology and evolutionary biology (e.g., research on population size distributions, trophic networks, and functional diversity) have shown that species interactions introduce stabilizing mechanisms, which can lead to deviations from strict power laws. For instance, Stephen Hubbell’s Unified Neutral Theory of Biodiversity and Biogeography suggests that species diversity and relative abundance within ecological communities can be explained by stochastic processes such as ecological drift and dispersal limitation, rather than by niche differentiation or fitness differences [64]. This theory assumes that trophically similar species are functionally equivalent, leading to species abundance distributions that can approximate power laws. Applying these principles to protein folds suggest that ecological interactions and stochastic processes can modulate their distributions, potentially shifting them toward PII-like distributions. These relationships help explain how ecological stability is influenced by factors such as species interactions and environmental perturbations, which in turn may affect observed abundance distributions.

### Fold abundance categories, geographic structuration of the repartition and eco-folds

This work also provide insights into the potential role of specific folds in the adaptive potential of planktonic species to abiotic variability they encounter along their transport by ocean currents. In particular, it showed that the distribution of particular folds in certain phyla exhibits a certain level of biogeographic structuring comparable to that of species communities. Three categories of folds have been identified according to the PII *k* parameter: rare, intermediate and abundant. In Chlorophyta, the biogeographic structuring of the abundant folds, which may function as *eco-folds* (biogeographic marker fold), besides being SFs for some of them, is consistent with that of the Mamiellophyceae communities. It is indeed known that their distribution is structured by biomes, basin as well as latitude, with Arctic communities being clearly distinct from the ones of the South Ocean, while Pacific communities are distinguished from those of the Atlantic [14], [65], [66]. It is therefore possible that temperature and iron, that are important drivers of the Chlorophyta distribution in the open Ocean [12], also partially drive the community-level biogeographic structure of the ones of at least some of the abundant folds in that phylum. Temperature is one of the primary factor impacting fold stability [67], while iron serve as an essential cofactor in many biological processes [68].

Furthermore, the three environmental categories of folds resemble the concept of unifolds, mesofolds and superfolds, which was initially developed based on genomic observations on collections of individual genomes [7], [8]. Since SFs exhibit a high tolerance to sequence changes, they can adapt to a wide range of biophysical conditions through sequence modifications. As a result, they are likely involved in several adaptive mechanisms. In contrast, unifolds, by definition, have a lower tolerance to sequence variability and perform highly specific functions. This limited tolerance to sequence variations may limit their adaptability to different physicochemical conditions, for example, by restricting their ability to modulate stability or flexibility of protein structure. Consequently, in environmental conditions that are specifically unfavourable to them, these folds could be replaced by alternative folds capable of performing the same function.

## Material and Methods

### Structural annotation of the proteomes

The structural annotation using CATH [21], [22] was performed on the proteomes of eukaryotic, prokaryotic and NCLDV MAGs of the *Tara* Oceans expedition (713 eukaryotes [53], 1,888 prokaryotes [53] available at https://www.genoscope.cns.fr/tara/#BAC_ARC_MAGs%20https://www.genoscope.cns.fr/tara/#SMA Gs and 30,802 NCLDV [54]) as well as on the 990 RPs from EukProt [55]. Fourteen eukaryote lineages, with a substantial number of MAGs and RPs, were selected for the study, some with slight differences in taxonomy between the MAGs and EukProt. Those lineages are the Choanoflagellata (11 MAGs « Choanozoa » that are all Acanthoecida and 27 RPs), the Arthropoda (219 MAGs and 19 RPs), the Chordates (44 MAGs and 22 RPs), the Chlorophyta (64 MAGs and 80 RPs), the Ciliates (30 MAGs and 45 RPs), the Haptophyta (92 MAGs and 27 RPs), the Cryptophytes (8 MAGs and 19 RPs), the MAST-4 (19 MAGs and 4 RPs), the Myzozoa (1 MAG MALV-I + 5 MAGs Myzozoa and 121 RPs), the Bigyra (13 MAGs and 3 RPs « Bicosoecida »), the Heterokontophytes (16 MAGs and 15 RPs « Core Oomycetes »), the Bacillariophyta (52 MAGs et 83 RP « Diatom ») and the Ochrophytes (51 MAG et 52 RPs « Chrysophyceae» « Eustigmatophyceae » « Pelagophyceae » « Dictyocophycea »).

The first step relies on scanning the Hidden Markov Models (HMMs) of Functional Family from CATH (funfam-hmm3.lib from 2020/09/22 [69], [70]) using the command *hmmsearch* from “*hmm-3.3.2”* [71] on the proteomes with a E-value threshold of 1e-10 (and the options -tblout and -noali). The HMM profiles are expected to represent the structural diversity of domains from major protein sequence databases, primarily Uniprot [72] and Ensembl [73]. The proteins on which the HMM scans are successful are then associated with a four digit code for each of their domains (one or several if the protein contains multiple domains).

This code reflects the classification of CATH into four hierarchical levels: Class (based on the proportion of α-helices and β-sheets), Architecture (describing the overall arrangement of secondary structures), Topology (reflecting fine structural similarities in terms of connectivity and spatial arrangement of secondary structures), and Homology (denoting high protein-sequence similarity, potential shared function, and close evolutionary relationships) [22]. Throughout this study, the CAT level is called “fold”.

### Fold occurrences

The OVs of Homologies (CAT**H**) in a proteome corresponds to the number of protein domains adopting it. From that value, the OVs of the folds (CA**T**) is the sum of all its CAT**H**, and so on up to the Class (**C**). The OV of each CATH at the level of a domain of life or a phylum is the sum of this CATH in every proteome that belongs to it. OVs were indeed calculated at different taxonomic and ecological scales: at the domain level (eukaryotes, prokaryotes, NCLDV), at the phylum level in both Eukaryotes and Bacteria and at the community level at a given sampling site (in that case, the OV of a CATH is the sum of all its OVs in every MAG present at the sampling site). OVs at the life domain level were displayed using the function *treemap* from the “*treemap*” R package [74](Supplementary Fig. 1A,3A).

The presence or absence of folds in MAGs, stations or phyla was assessed using the OVs, transforming any value greater than one by one. This dataset is referred to as the fold *repertoire* in the following analysis. The number of combinations of each fold is the number of different mutidomain proteins the fold is found in; the number of partner was determined by counting how many different folds co-occurred with it in multidomain proteins.

### Models of fold occurrence distributions in genomes

The fold OVs in the MAGs, RPs and marine planktonic communities can be analysed by examining their mathematical distribution (*i.e*. the number of folds exhibiting a specific OV). In order to reduce statistical noise in the data [54], logarithmic binning was applied using the *logspace* function from the “*ramify”* R package [74].

OVs were first transformed into relative values to allow direct comparisons between MAGs and RPs, between MAGs from the different domains of life and phyla within Eukaryotes and Bacteria, and between communities. These transformed values were grouped into bins on their average relative occurrence value, with bin intervals regularly spaced on a logarithmic scale. The power-law function:

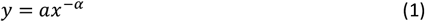

was used to perform the fits using the *lm* function from base R [75](Supplementary Fig. 1B, 2, 3B, 4). The power-law models on the OV distributions were displayed using the “*ggplot2*” R package with a double log10 scale [76].

### Fold abundances in marine planktonic communities

To study the biogeography of folds, their abundances in the environment were calculated. An AV was calculated for each identified CATH ID by taking into account both the relative abundance of each species [53] in the targeted station and phylum, as well as the OVs of these CATH ID in proteomes. This measure corresponds to the sum of the occurrences of a given fold, weighted by the vertical metagenomic sequence coverage of each species encoding it. AVs were calculated using the following formula:

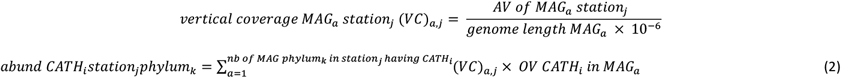

Which can be generalized for all CATH ID and stations by phylum, with *m* representing the number of MAGs in phylum *k* and *n* denoting the size of the fold repertoire in phylum *k*, as follows :

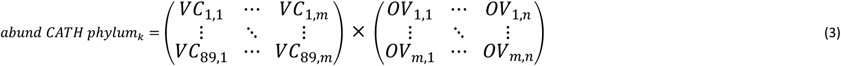

And for the whole eukaryotic and bacterial communities with:

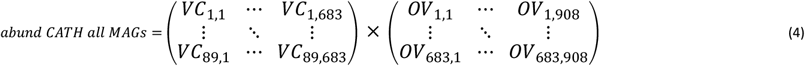

We accounted for the individual incompleteness of each MAG by using their vertical coverage. The abundance of each Topology was calculated as the sum of all the Homologies adopting that specific Topology.

Since the relative abundance of the MAGs in a given metagenome is compositional, a Centered-Log Ratio (CLR) transformation was applied to the fold AVs to mitigate certain biases associated with compositional data [77], [78], [79]. To address the numerous zeros in the dataset, the *decostand* function with the “rclr” method from the “*vegan*” R package [80] was used. Regarding eukaryotic data, biogeographic analyses were performed on surface samples of the 0.8-2000µm size fraction [9], [81]. To represent fold abundances on a heatmap, a distance matrix was computed using the *vegdist* function with the “bray” method, then clustered with the *hclust* function with the “average” method, both from the “*vegan*” R package [80] (Fig. 1A; Supplementary Fig. 5A, 13). The same method was used with the eukaryotic MAGs (Supplementary Fig. 13)

Similarities between stations in terms of fold abundance were estimated using Principal Coordinates Analysis (PCoA) with the *cmdscdale* function (parameters: k=3, eig=T and add=T) from base R [75], applied to a distance matrix produced using *vegdist* with “*bray*” method [80]. The sampling sites coordinates on the three first PCoA axes were then converted into Red-Green-Blue (RGB) color codes following the same principle as in Richter *et al*. [11], using the *rgb* function from base R [75]. These colors were mapped onto a world map using the “*world*” map from the “mapdata” R package [82](Fig.2 B) (Fig. 1B). The distribution of community OVs and AVs was first fitted to a linear model. To ensure comparability between AVs and OVs, and given that the CLR transformation alters the shape of the distribution, AVs were only transformed to relative values (as was done for OVs). Both distributions were then log-binned into 100 bins. The model used for the fits follows the equation (1), and the same methodology as described in “linear models of fold occurrence distributions in genomes” was applied (Fig. 2A). The fits were calculated with the *linregress* function from the “*SciPy*” Python package [83] and plotted using *matplotlib* [84].

### Pareto Type II fits on the distribution of fold abundances

After performing the linear fit described above, the Probability Density Function (PDF) of the PII law (also known as the Lomax distribution [85], [86]):

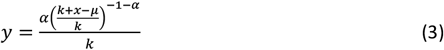

was selected to model the distribution of relative AVs of folds in eukaryotic and bacterial communities.

PII fits were performed using a custom Python script that includes the *curve_fit* function from the “*scipy.optimize*” package [83] with default parameters except for the boundaries, which were set between 0 and the maximum occurrence for both *k* and *mu*, and with *maxfev* set to 50,000. A range of alpha values between 0 and 4.1 was tested and the Root Mean Square Error (RMSE) and the variance of the parameter estimate were used to choose the best combination of parameters.

The model with the lowest variance value within the first quartile range of RMSE values was selected as the best fit. If the *k* value from the best fit is less than 10 times larger than the lowest occurrence value, typically occurring when low occurrence values contain outliers, the script adjusts the data by removing the lowest value and repeats the fitting process.

Once the optimal parameter combination for the PII distribution is identified, a KS test is performed using the *ks_2samp* function from the “*scipy*.stats” Python package [83], to assess the goodness-of-fit between the model and the observed data. The resulting fits were plotted using *matplotlib* [84] (Fig. 2B; Supplementary Fig. 6).

The results of the KS test (KS value and *p*-value) were plotted on a world map for each phylum using the “*world*” map from the “*mapdata*” R package [82] using a significance threshold set at 0.01 (Fig.2 C, Supplementary Fig.7). The density distributions of both KS statistics and *p*-values were visualized per phylum and for all eukaryotes using the *geom_density_ridges* function from the “*ggridges*” R package [87] (Supplementary Fig.8 A). Correlations with environmental and non-environmental parameters were tested using the *cor* function from base R [75] with defaults parameters, as well as the *cor.mtest* function from the “corrplot” package [88], with the parameter of confidence level *conf.level* set to 0.99. The results were displayed using the *ggcorrplot* function from the “ggcorrplot” package [89](Supplementary Fig. 8B-C). Most non-environmental parameters, including BUSCO completion, genome length and coding length, were published alongside the MAGs [53]. The Shannon index of the MAGs per phylum was computed using the *diversity* function with default parameters from the “*vegan*” R package [80]. The environmental parameters selected for this analysis were SunShine Duration (SSD), temperature at the time of sampling, salinity, chlorophyll-*a* concentration, dioxygen concentration and nitrogen concentration in three different forms (NH_4_, NO_3_ and NO_2_). Additionally, iron, phosphate and silicate concentrations were included [9], along with the median, amplitude and standard deviation of monthly temperature [56]. The only parameters that were not measured *in situ* during the TO expedition were iron and phosphate concentrations, variables related to monthly temperatures and salinity. Iron concentrations on a global scale were provided from the biogeochemical model PISCES-v2 [15], [57]. Monthly temperatures, later transformed into annual median and standard deviation values, as well as phosphate concentrations and salinity, were extracted from WOA13 [56]. Frémont [15] reported a strong correlation between *in situ* TO measurement and values obtained from WOA13 (*r*^2^ = 0.96 for temperature, 0.89 for phosphate concentration and 0.83 for salinity). Missing values from WOA13 and PISCES-v2 at TO sampling stations were extrapolated by assigning the value from the closest available longitude.

Finally, the values of the parameter *k* were used as thresholds to classify folds based on their relative abundance across stations within each phylum. This led to the creation of three non-core categories in each phylum. The first category corresponds to folds with abundances lower than *k* in all samples; the second category correspond to folds with abundances always above *k* in all samples. Folds that do not consistently fall above or below *k*, varying in abundance across different samples, were classified the “intermediate” category. The core of each category was then established by selecting the folds that belong to the same category in the six selected eukaryotic phyla. Folds classified as first category in every phylum were assigned to the core first category, and similarly for the second and intermediate categories (Supplementary Fig. 9).

The properties of the folds, including the number of folds, their Architecture, and their number of partners, were analysed for each of the core and non-core categories. These properties were displayed using the *upset* function from the “*ComplexUpset*” R package [90], [91] as well as functions from “*ggplot2”* (Supplementary Fig. 10, 11). Significant differences in fold versatility among the different categories within each phylum were tested using the *geom_signif* function from the “ggsignif” package with the Wilcoxon test with a 99% confidence threshold(Supplementary Fig. 13).

The average membership of each fold in the three environmental categories across the six phyla was established using different classification criteria: the designation “in most phyla” indicates that a fold belongs to a given category in at least four phyla. The term “when present” refers to cases where some folds are absent from the fold repertoire of some phyla. Finally, “absent from the selected phyla” indicate that folds are not found in the repertoire of any of the six phyla selected for the analysis (Supplementary Fig. 14).

## Structuring of the biogeographic distribution of the folds from each abundance category

The AVs of the folds of each core category were summed to create a seventh “phylum”, grouping the six phyla together, and referred to as “6 phyla together” in the following analysis.

Differences in AVs between the fold of the categories defined above were tested within each phylum using a Wilcoxon test with a 99% confidence threshold with the *geom_signif* function (Fig. 3A; Supplementary Fig. 13).

The AV variability of folds from each category and phylum across communities was first examined using α-diversity. AVs were CLR-transformed and translated as described in the “fold abundances in marine planktonic communities” section. The α-diversity was then computed using the Shannon index, following the same method as described for the MAGs in the “Pareto Type II fits on the distribution of fold abundances”. The results were plotted using the *geom_smooth* function of the “*ggplot2*” R package [76] with default parameters (Fig. 3B; Supplementary Fig. 16A). The difference of Shannon index between polar and non-polar stations was then estimated. Polar stations were defined as those with absolute latitudes above 60°, and non-polar stations as those below 60°. The significance of the differences was tested for each phylum and category using a Wilcoxon test, following the methodology described in previous sections (Supplementary Fig. 15, 16B; Supplementary Table 2). The same analysis was done with the AVs of the MAGs (Supplementary Fig. 17; Supplementary Table 2).

The structuring of the biogeographic distribution of folds within each category and phylum was then assessed using a dbRDA. The environmental dataset for the model consisted of a matrix containing both environmental parameters and Moran Eigenvectors Maps (MEMs). MEMs were calculated with the *dbmem* function with MEM.autocor parameter set to “non-null” from the “*adespatial*” R package using the in-water distances between TO stations [14]. To retain only the most relevant Moran Eignen Vector Maps (MEMs) for the dbRDA analysis, the *ordistep* function from the “*vegan*” R package [80] was used with the direction parameter set to “both” and 10,000 permutations were performed on both a null model and a model incorporating all MEMs as explanatory variables. Only MEMs with a *p*-value below 0.01 were retained. The environmental parameters used for this analysis are from the same source as those used in “Pareto type II fits on the distribution of fold abundances”. Only temperature, temperature amplitude, iron and phosphate concentrations were retained for this analysis. Statistical significance of these parameters was tested using the function *anova* from the “*car”* R package [94] with parameters *by* set to “margin” and 999 permutations. A distance matrix was then calculated on the rclr-transformed AVs using the *dist* function from base R [88] with default parameters. Subsequently, a Principal Coordinates Analyses (PCoA) was performed using the *pcoa* function from the “*ape*” R package [95] with default parameters. The results of that PCoA were then used as input for the subsequent analysis. Ultimately, the dbRDA was calculated with the results of the PCoA and the selected MEMs plus environmental variables as explanatory factors. The adjusted determination coefficient of the model was estimated using the *RsquareAdj* function from the “*vegan*” R package [80]. The dbRDA analysis was also performed using the AVs of the MAGs per phylum with the same environmental and MEM data. The coordinates of the stations in the two first dimension of the dbRDA space were translated into a RG code, following the same procedure as in “Pareto Type II fits on the distribution of fold abundances”. The TO stations were then displayed on a world map and coloured according to that code (Fig. 3C; Supplementary Fig. 18, 19, 20; Supplementary Table 3-4-5).

## Acknowledgements & Fundings

LP was funded by the Doctoral School “Structure and Dynamics of Living Systems” of UEVE/Université Paris-Saclay. AL and LJ were funded by the Université d’Évry (UEVE). CV received financial support from ANR SeqDigger (ANR-19-CE45-0008). JR received financial support from ANR CLIMACLOCK (ANR-20-CE20-0024). EP acknowledges a fellowship funded by the Stazione Zoologica Anton Dohrn (SZN) within the SZN-Open University Ph.D. program and the European Union’s Horizon 2020 Blue Growth research and innovation programme under grant agreement no. 862923 (project AtlantECO). It benefited from access to high-performance computing resources through GENCI-[TGCC/CINES/IDRIS]. Computation for structural protein annotation was done on the OSU Pythéas cluster with the help of C. Blanpain, J. Lecubin and SIP members. We thank the commitment of the Research Federation for the Study of Global Ocean Systems Ecology and Evolution (FR2022/TaraGOSEE) and of Stazione Zoologica Anton Dohrn. We thank LAGE (Laboratoire d’Analyses Génomiques des Eucaryotes, CEA) members for stimulating discussions on this project, C. Scarpelli and members of the scientific computation team from Genoscope for support on computations. We thank all members of the *Tara* Oceans consortium for maintaining a creative environment and for their constructive criticism. *Tara* Oceans would not exist without the *Tara* Ocean Foundation and the continuous support of 23 institutes (https://oceans.taraexpeditions.org/). This article is contribution number XXX of *Tara* Oceans.

## Author contributions

LP, ML, YT and OJ conceptualized the study and designed the research. LP, CV, JR, EP and ML conceptualized and developed the methodology. LP, AL, LJ and EP implemented the methodology. LP, EP, DI, ML, YT and OJ interpreted the results. LP, ML, YT and OJ wrote the manuscript; JR, EP and DI contributed to writing. All authors have read and approved the manuscript.

