## supplementary figures for "Deviation from Power-Law Distribution when Scaling the Distribution of Marine Plankton Folds from Genomes to Communities"

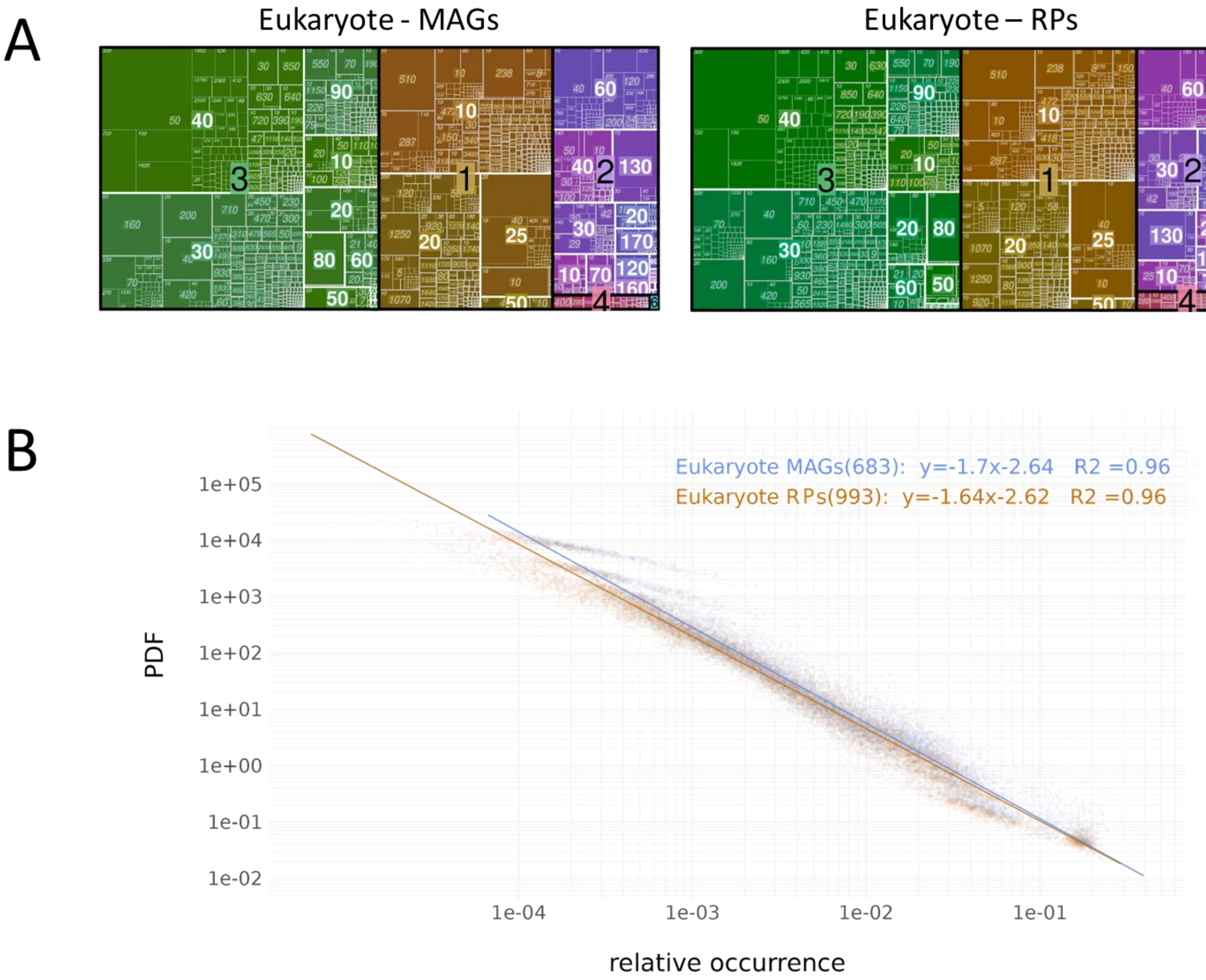


**Supplementary Figure 1. Comparisons between the eukaryotic environmental genomes (MAGs) and reference proteomes (RPs) in terms of fold composition. (A)** Treemaps for the MAGs (left) and the RPs (right). The size of each square represents the proportion of occurrences of a given CATH ID when combining all the genomes of the domain from each database. The size of each number corresponds to a hierarchical level. From largest to smallest: Class, Architecture, Topology, Homology. Each colour shade represents one Class: brown = Class 1; purple = Class 2; green = Class 3; red = Class 4; blue = Class 6. **(B)** Power law models of the distribution of the relative occurrence values (OVs) of the folds in MAGs (blue) and RPs (orange). The relative OVs and the probability density function (PDF) of each bin of OV are shown on the x-axis and y-axis respectively. Both axis are on a log10 scale. Each dot represents a bin with a given OV in one proteome. The grey zone surrounding the lines indicates the standard error. In the top right corner from left to right: dataset name and number of proteomes it contains, regression equation, and adjusted $R^2$ value. Every fit is statistically significant at a 99% confidence threshold.

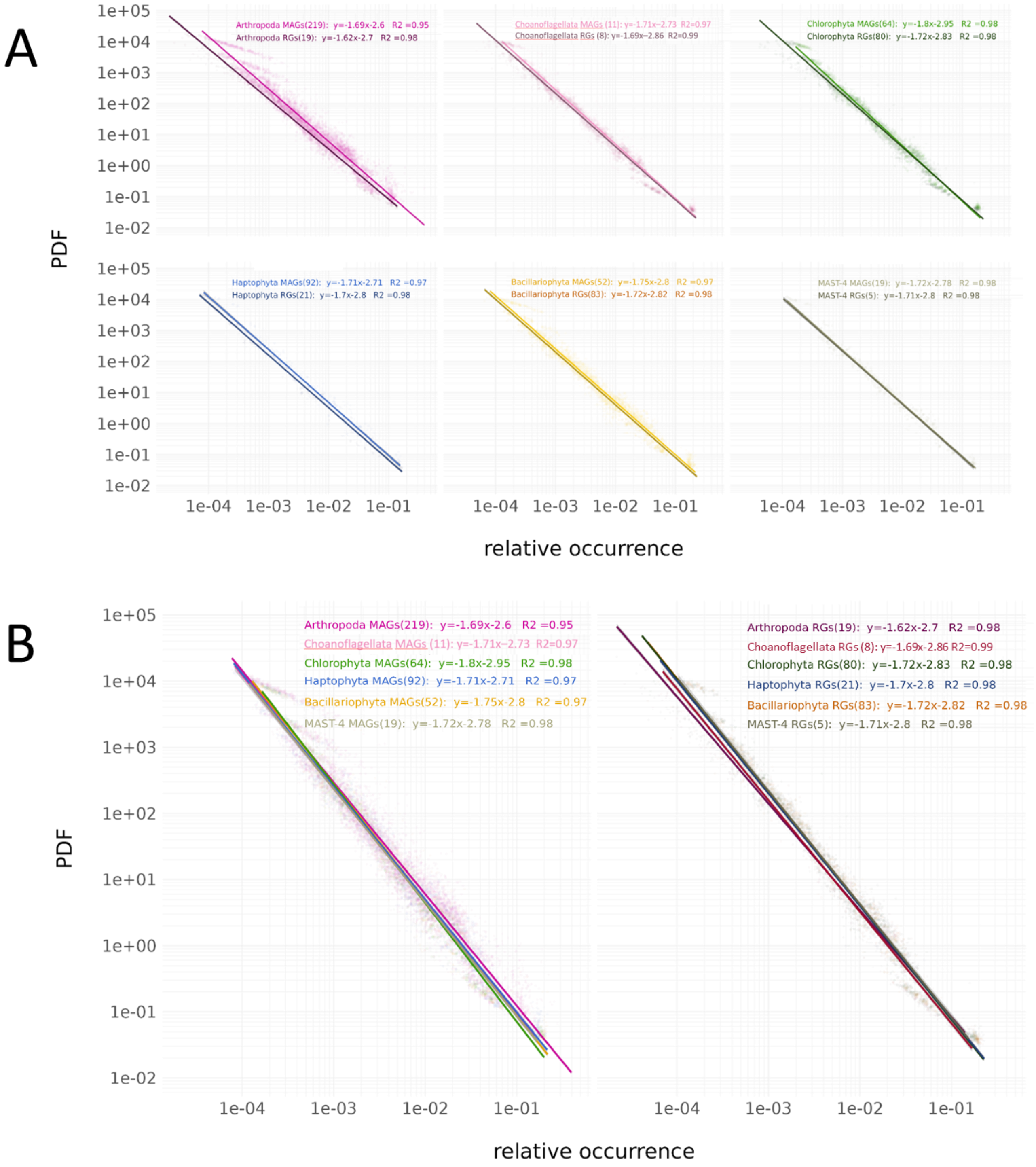


**Supplementary Figure 2. Comparison of the power law models at the phylum scale between eukaryotic MAGs and RPs.** Power law models of the distribution of the relative OVs of the folds in. The relative OVs and PDF of each bin of OV are shown on the x-axis and y-axis respectively. Both axis are on a log10 scale. Each dot represents a bin with a given OV in one proteome. The grey zone surrounding the lines indicate the standard error. In the top right corner from left to right: dataset name and number of proteomes in contains, regression equation, adjusted $R^2$ value. Every fit is statistically significant at a 99% confidence threshold. **(A)** Pairwise comparisons of the power law models per phylum (from left to right and top to bottom: Arthropoda, Choanoflagellata, Chlorophyta, Haptophyta, Bacillariophyta, MAST-4). **(B)** Comparison of the power law models per database (left: MAGs; right: RPs).

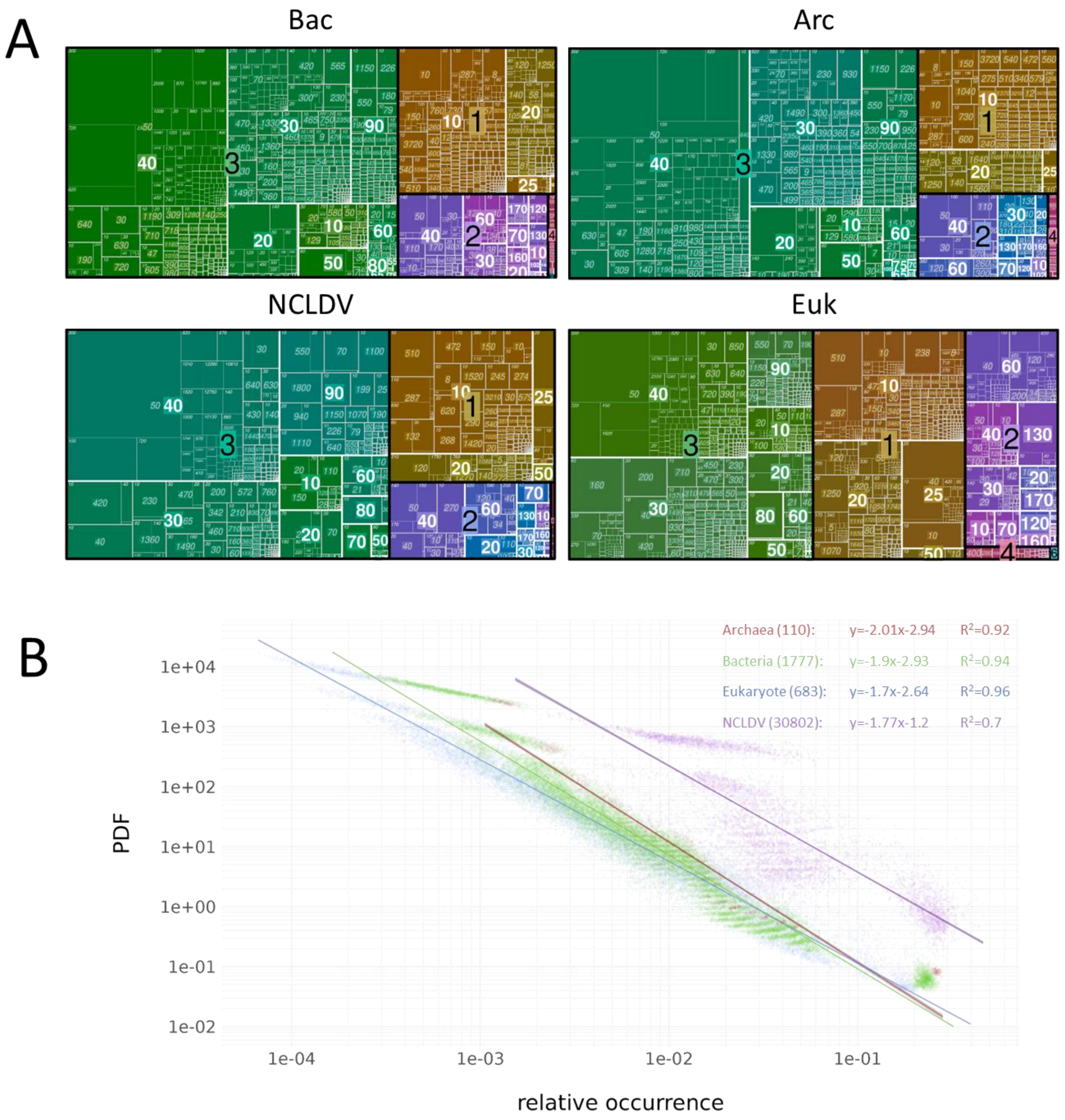


**Supplementary Figure 3. Distribution of folds in the MAGs from the different domains of life. (A)** Treemaps for the Bacteria (Bac), Archaea (Arc), NCLDV and Eukaryotes (Euk). The size of each square represents the proportion of occurrences of a given CATH ID when combining all the genomes of the domain from each life domain. The size of each number corresponds to a hierarchical level. From largest to smallest: Class, Architecture, Topology, Homology. Each colour shade represents one Class: brown, Class 1; purple, Class 2; green, Class 3; red, Class 4; blue, Class 6. **(B)** Power law models of the distribution of the relative OVs of the folds in Archaea (red), Bacteria (green), Eukaryotes (Blue) and NCLDVs (purple). The relative OVs and PDF of each bin of OV are shown on the x-axis and y-axis respectively. Both axis are on a log10 scale. Each dot represents a bin with a given OV in one proteome. The grey zone surrounding the lines indicate the standard error. In the top right corner from left to right: dataset name and number of proteomes in contains, regression equation, adjusted $R^2$ value. Every fit is statistically significant at a 99% confidence threshold.

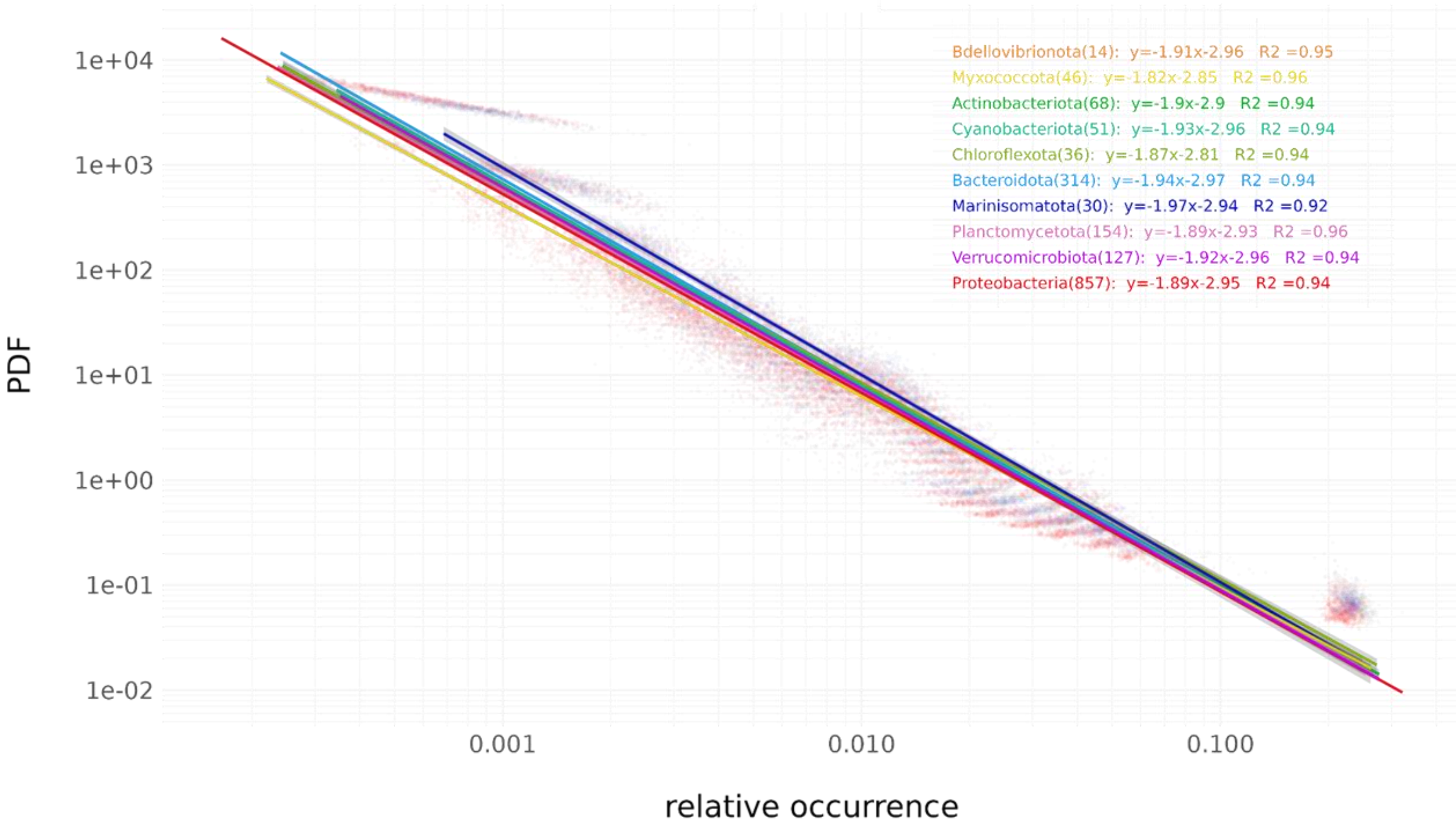


**Supplementary Figure 4. Power law models of the distribution of folds in bacterial phyla.** Power law models of the distribution of the relative OVs of the folds in different bacterial. The relative OVs and PDF of each bin of OV are shown on the x-axis and y-axis respectively. Both axis are on a log10 scale. Each dot represents a bin with a given OV in one proteome. The grey zone surrounding the lines indicate the standard error. In the top right corner from left to right: dataset name and number of proteomes in contains, regression equation, adjusted $R^2$ value. Every fit is statistically significant at a 99% confidence threshold.

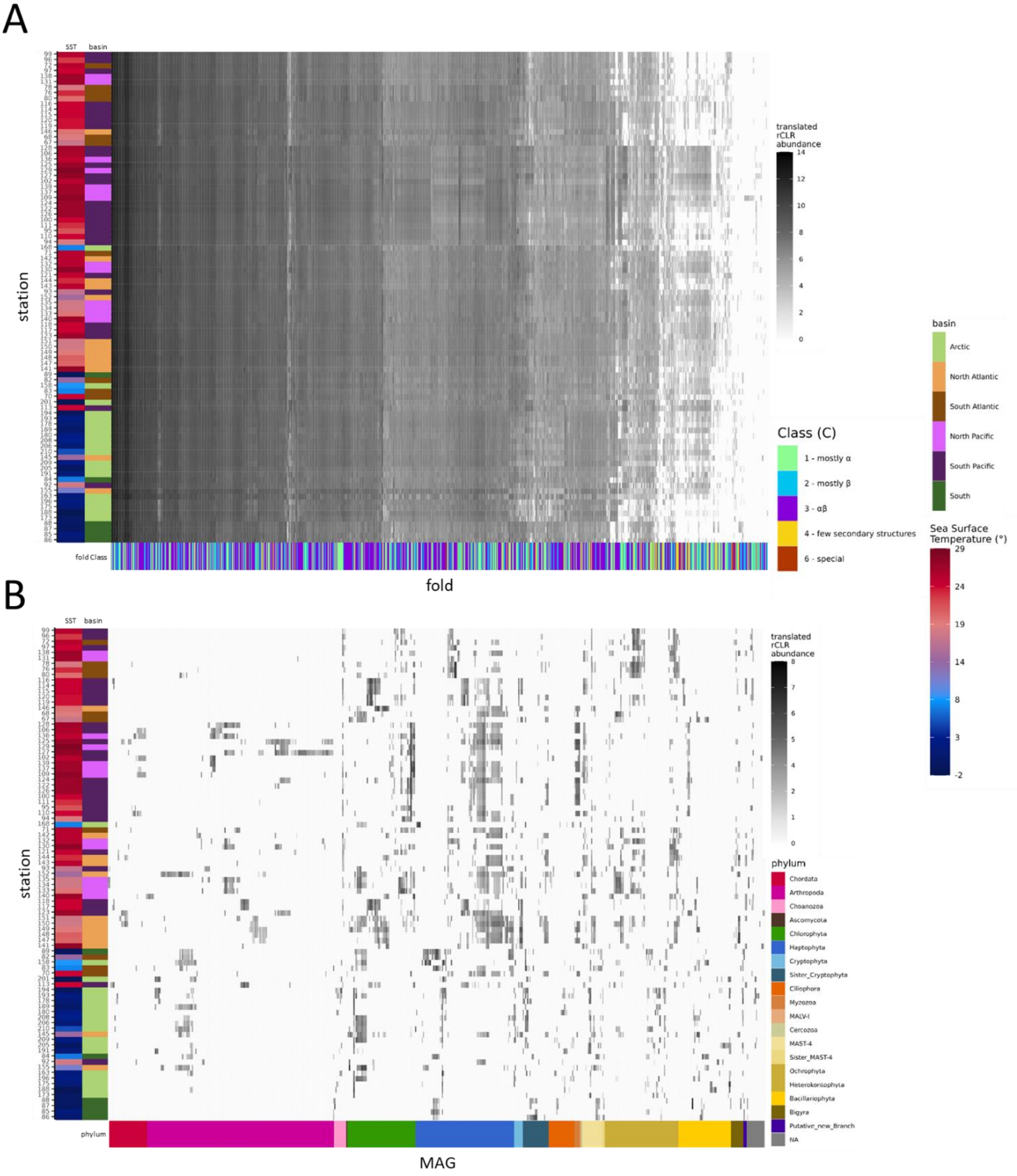


**Supplementary Figure 5. Abundance of folds and MAGs of the 0.8-2000µm size fraction across 89 *Tara* Oceans (TO) surface stations. (A-B)** The clustering on the y-axis is the same as the one in Fig. 1A. **(A)** Identical to Fig. 1A, but with additional layers on the y-axis (Sea Surface Temperature (SST), basin), and on the x-axis (fold Class). **(B)** Heatmap of MAG abundance values (AVs), with MAGs displayed along the x-axis. They were first grouped by phylum (represented by the additional coloured layer under the x-axis) and subsequently clustered within each phylum based on their AVs across the stations. Stations on the y-axis follow on the same order as in **(A)** and Fig. 1A.

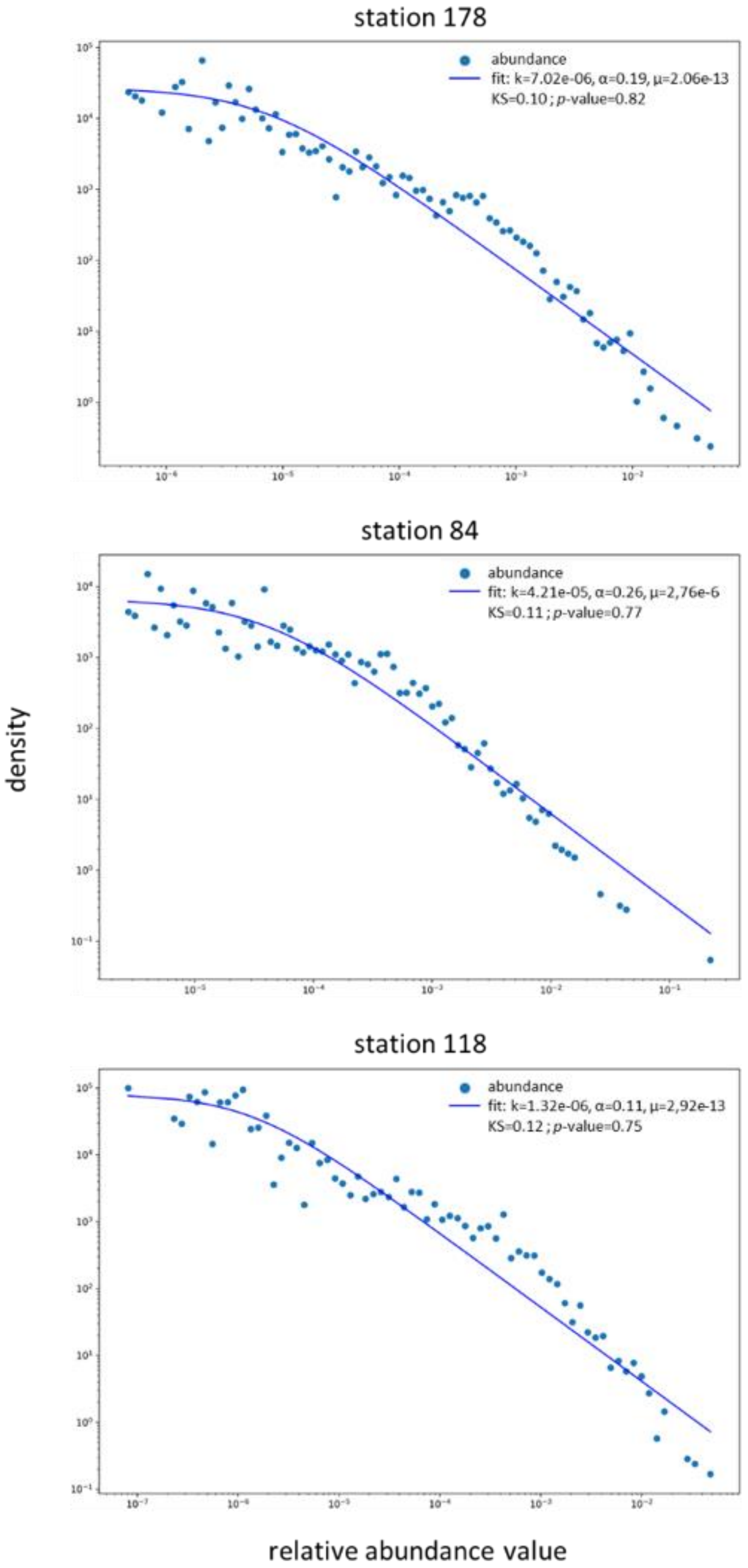


**Supplementary Figure 6. Examples of Pareto type II (PII) models on the distribution of fold AVs of bacterial communities.** From top to bottom: station 178, 84 and 118. The values of the equation parameters (k, α, μ), the Kolmogorov-Smirnov (KS) test and its associated *p*-value are on the top right corner of each fit.

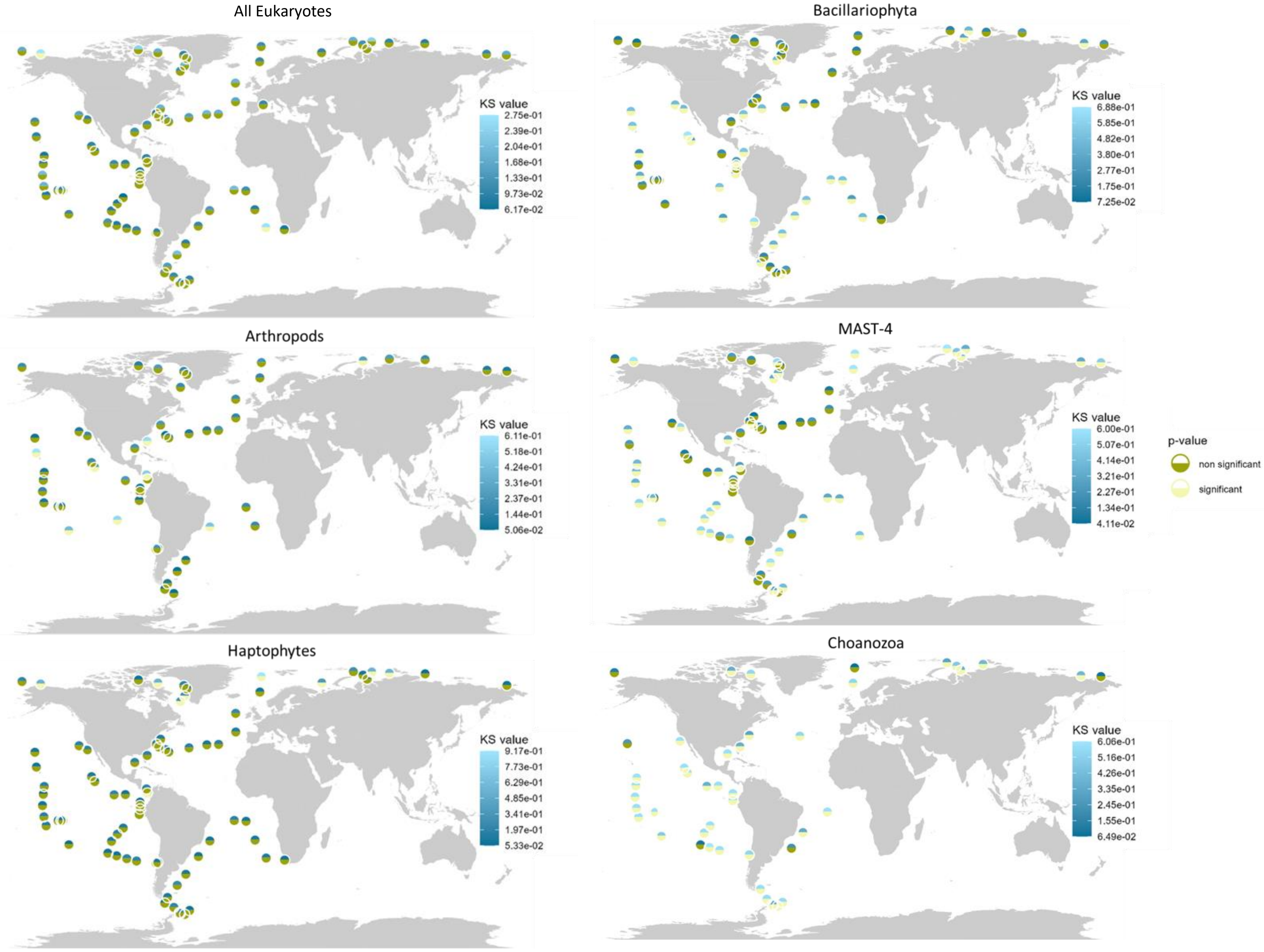


**Supplementary Figure 7. KS and *p*-value of the PII models in the TO stations in different eukaryotic phyla.** Each dot represents a TO station. The upper colour in shades of blue corresponds to the KS test value (the lower, the better the fit); the lower, in shades of green, represents the *p*-value of the test where the null hypothesis $H_0$ is *the distribution of the observed data follow the Pareto type II distribution of given parameters*. In stations where $H_0$ cannot be rejected (log10(*p*-value) > -2), the data are accordingly considered to follow the Pareto type II distribution.

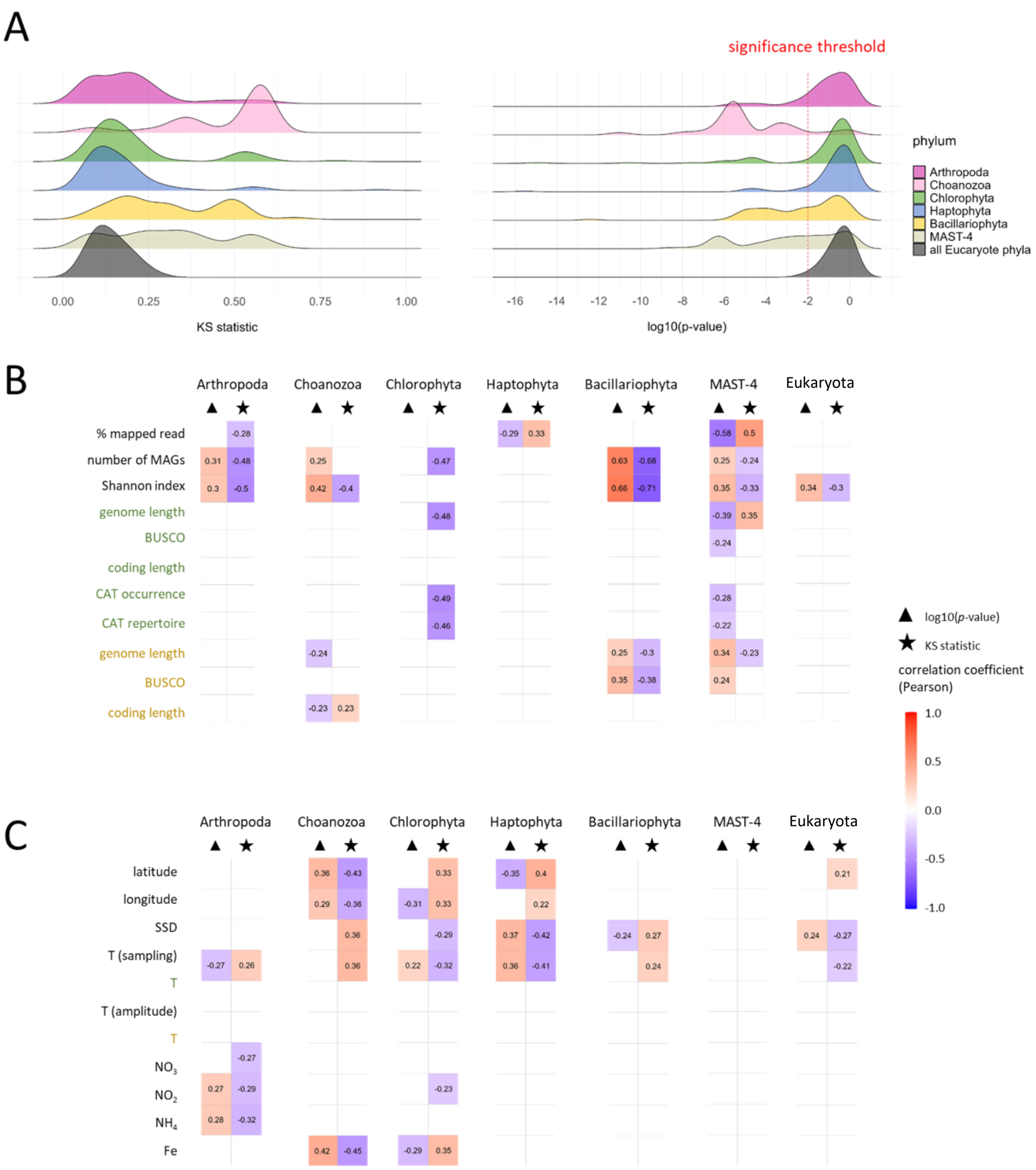


**Supplementary Figure 8. Variability of the quality of PII fits across stations and eukaryotic phyla. (A)** Distribution of densities of the KS test values (left) and the associated *p*-value with the null hypothesis $H_0$: *the observed data follow the Pareto type II distribution of given parameters*. A vertical red dotted line in the plot indicates the significance threshold (1%). The lower the KS value, the better the fit (leftmost part of the plot on the left). **(B-C)** Triangle and stars on top of each plot correspond to the log10 of the *p*-value and the KS statistic, respectively. Names in green and yellow on the y-axis correspond to the mean and standard deviation of the variable values, respectively. “Eukaryota” corresponds to the data obtained with all eukaryotic MAGs. **(B)** Correlation between KS value, log10(*p*-value) and non-environmental parameters associated with each TO station for each eukaryote phylum and “Eukaryota”. CAT repertoire: number of different CAT in each genome. CAT occurrence: sum of the occurrence of all CAT in each genome. **(C)** Correlation between KS value, log10(*p*-value) and environmental parameters associated with each TO station for each eukaryote phylum and “Eukaryota”. SSD: sunshine duration. T(sampling): sea surface temperature (SST) measured at the time of the sampling [1]. T: SST coming from World Ocean Atlas (WOA [2]). T(amplitude): difference between maximal annual SST and minimal annual SST in WOA. NO2, NO3, NH4: nitrite, nitrate and ammonium concentration measured during the TO expedition. Fe: iron concentration estimated by PISCES-V.2 [3], [4].

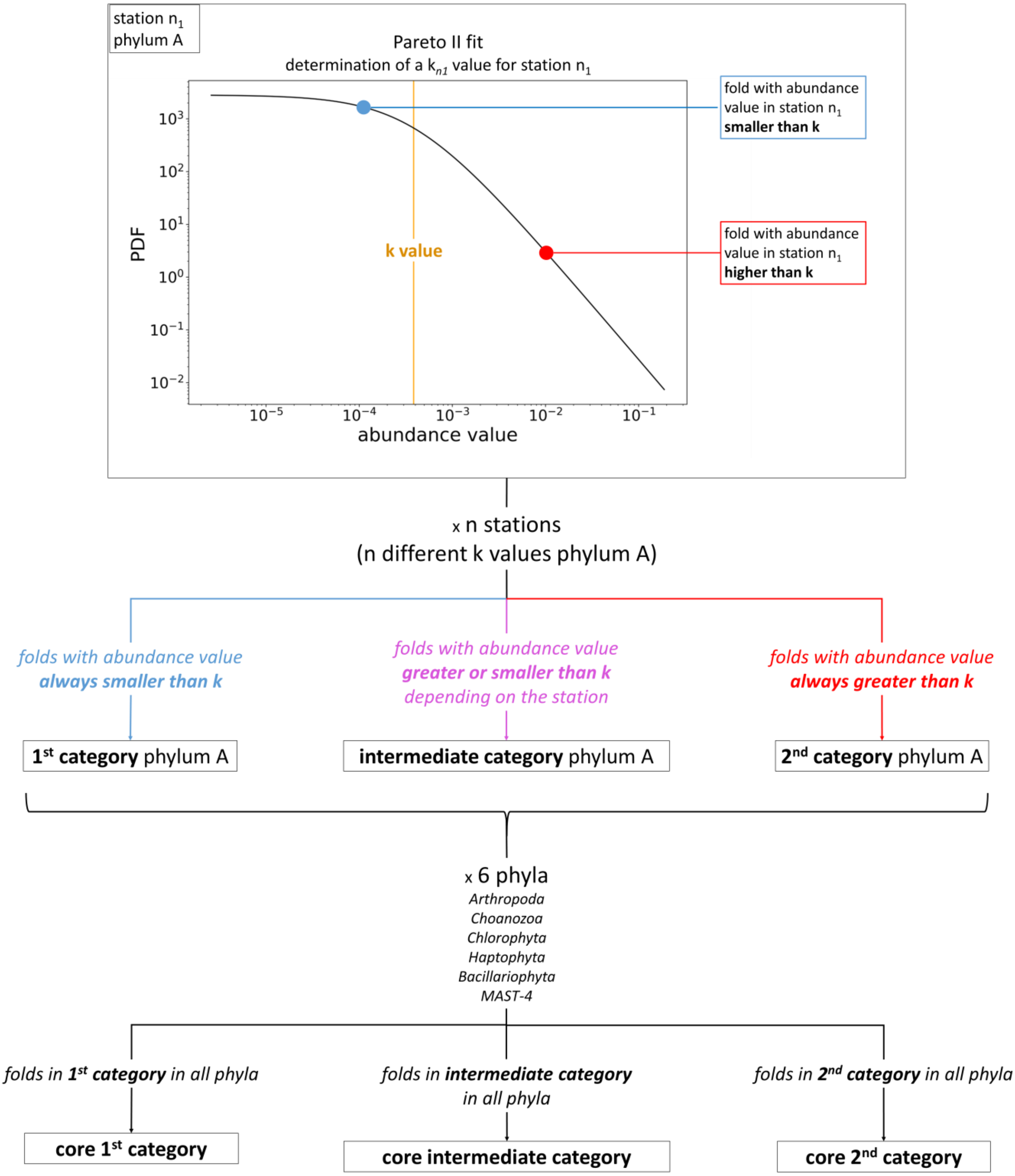


**Supplementary Figure 9. Schematic definition of the abundance fold categories.** From top to bottom, the workflow starts with the PII model of the AV distribution in station $n_1$ and phylum A. Blue boxes and texts correspond to the cases where the fold AV is lower than *k*. Red boxes and texts correspond to the cases where the fold AV is greater than *k*. The purple boxes and texts indicate the intermediate situation. The same approach is performed for the six eukaryote phyla selected in this part of the study. Folds belonging to the same category in all phyla are annotated as « core ».

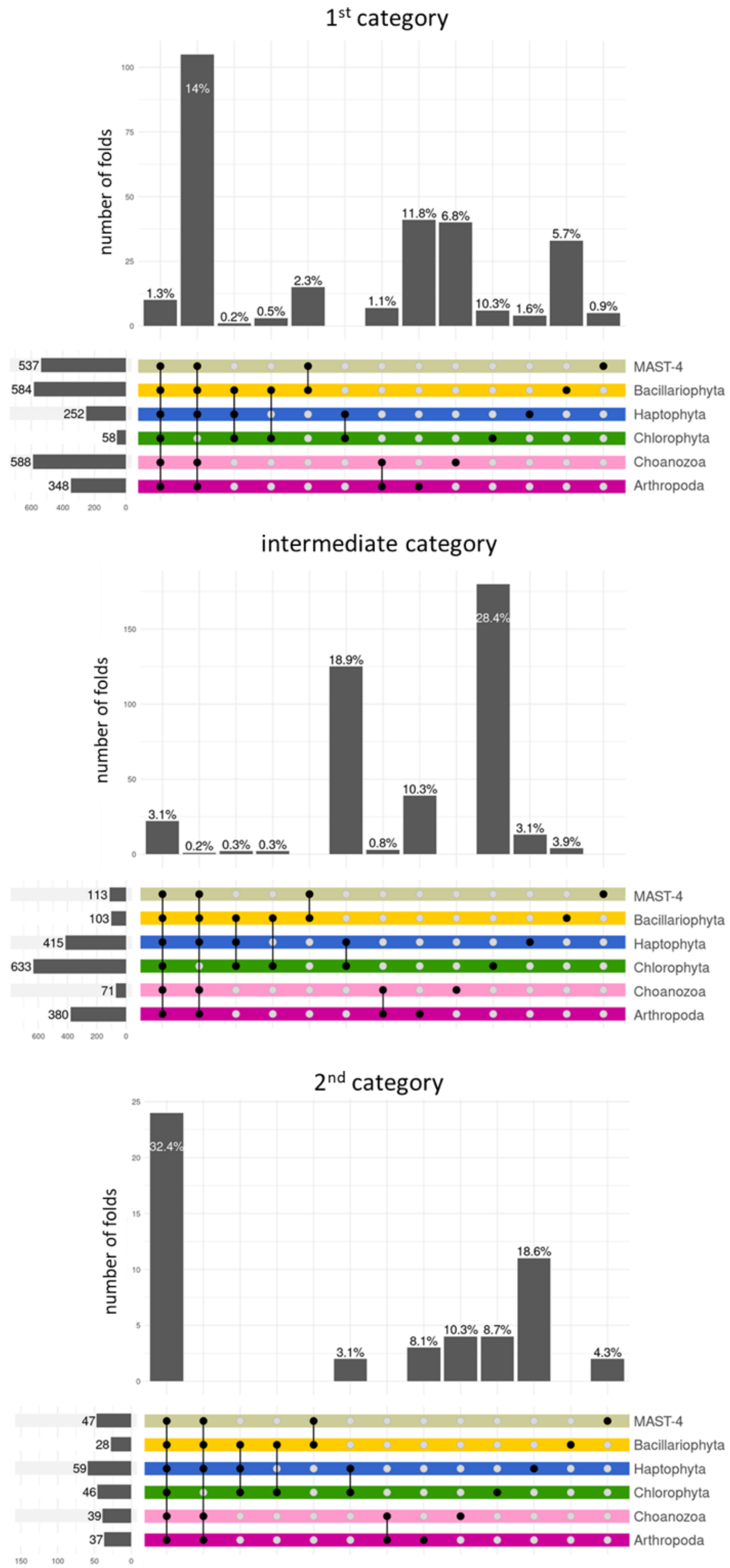


**Supplementary Figure 10. Number and proportion of folds in each abundance category of fold and eukaryote phyla.** Upset plots representing the intersections of phyla per category (top: first category; middle: intermediate; bottom: second category). The vertical bars above the coloured horizontal lines show the number of folds in each intersection. The number on top of each bar indicates the proportion of folds related to the total fold diversity in that category, all phyla together. The horizontal bars on the left correspond to the number of folds in each category and phylum.

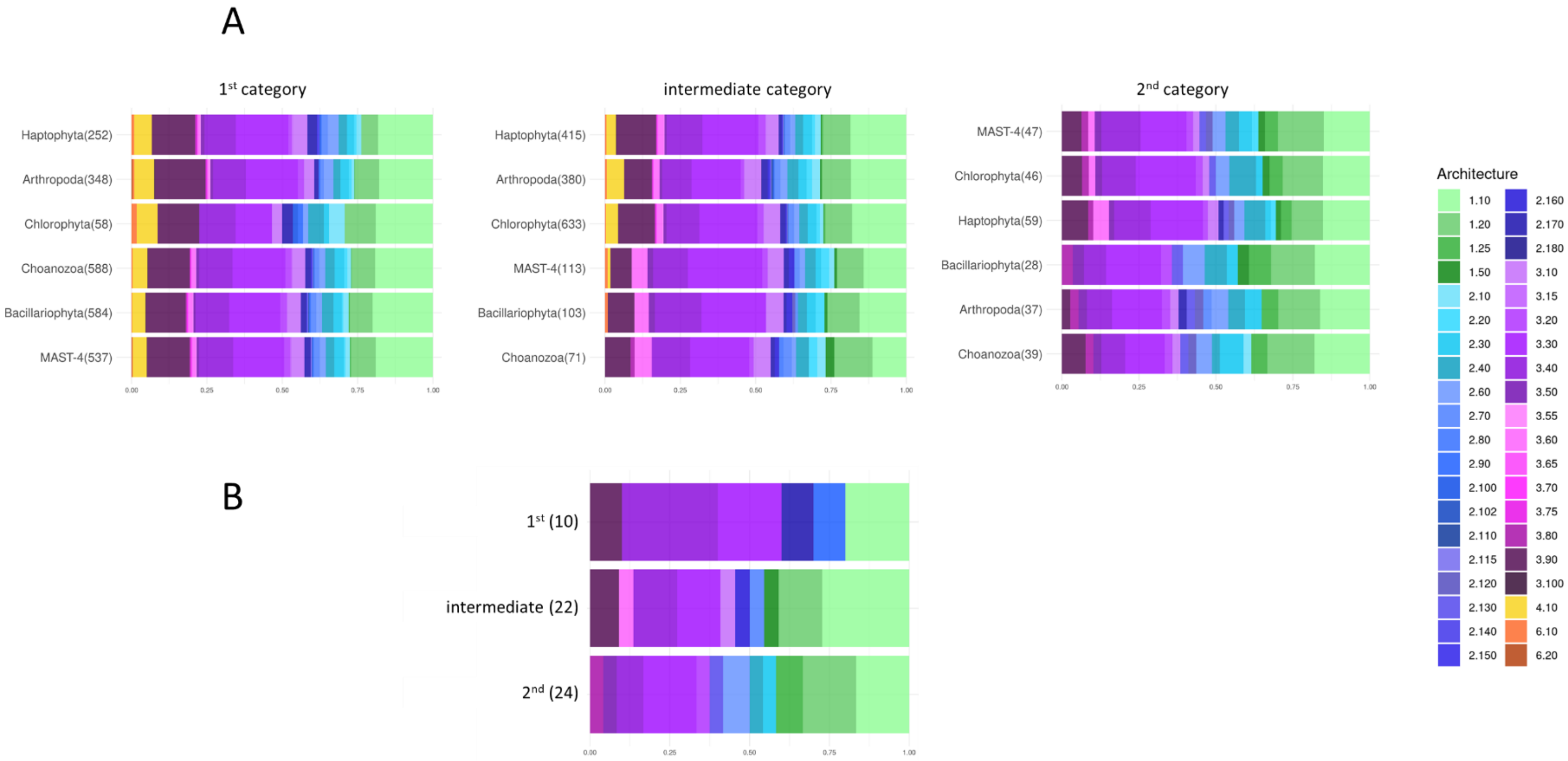


**Supplementary Figure 11. Number of folds and composition in Architecture of the environmental categories (core and non-core per eukaryote phylum). (A-B)** Barplots of the composition in Architecture of each non-core category per phylum **(A)** and for the core **(B)**. The number of folds in each category and phylum is indicated in brackets after the phylum or the category name. The x-axis represents the proportion of folds belonging to each Architecture in the categories and phylum. The colours correspond to the Architectures. In shades of green: Architectures of Class 1. In shades of blue: Architectures of Class 2. In shades of pink: Architectures of Class 3. In yellow the Architecture 4.10. In shades of orange the two Architectures of Class 6. **(A)** The phylum are ordered on the y-axis by similarity in terms of composition in Architectures.

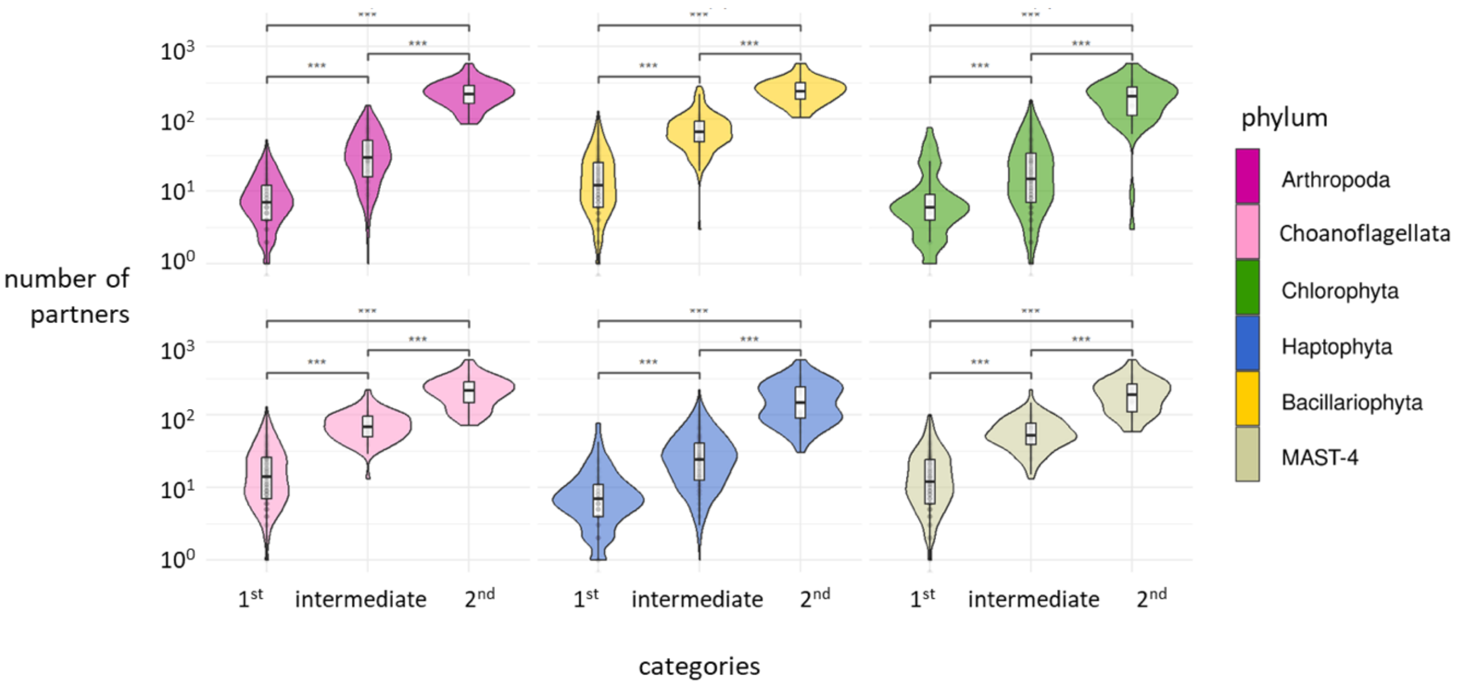


**Supplementary Figure 12. Number of partners of each fold per fold abundance category and eukaryote phylum.** The fold categories are indicated on the x-axis. The number of partners (number of different folds in combination with each fold) is indicated on the y-axis. Statistical significance was tested with a Wilcoxon test (***: *p*-value < 0.01).

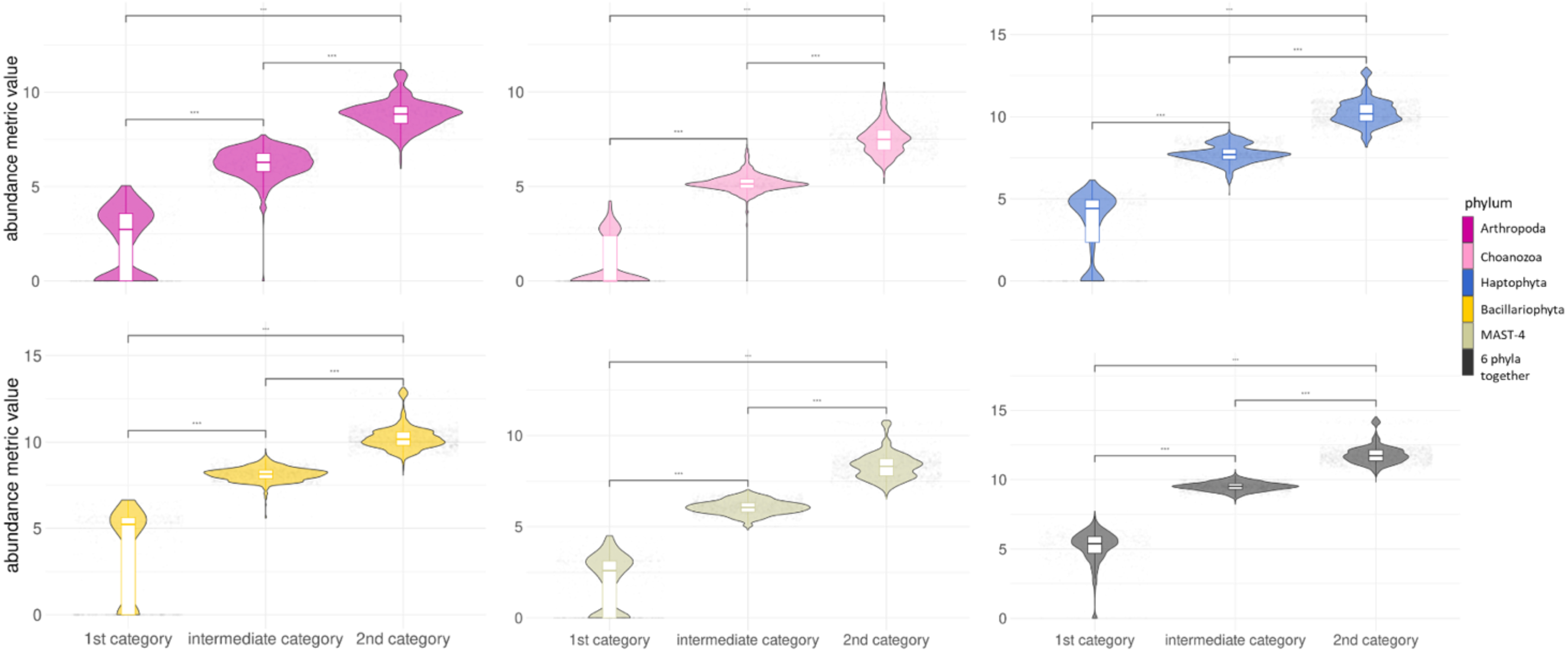


**Supplementary Figure 13. Abundance differences between fold abundance categories and eukaryote phylum.** The categories are indicated in the x-axis of each facet (from left to right: first, intermediate, second). From top to bottom and left to right: Arthropoda, Choanoflagellata, Haptophyta, Bacillariophyta, MAST-4 and the six phyla together. The results for the Chlorophyta are displayed in Fig. 3A. The CLR translated AVs of the folds are indicated on the y-axis. Statistical significance was tested with a Wilcoxon test (***: *p*-value < 0.01).

| core category | CAT id | Class | Architecture | Topology |
|---|---|---|---|---|
| 2nd | 1.10.10 | Mainly Alpha | Orthogonal Bundle | Arc Repressor Mutant, subunit A |
| 2nd | 1.10.287 | Mainly Alpha | Orthogonal Bundle | Helix Hairpins |
| 2nd | 1.10.510 | Mainly Alpha | Orthogonal Bundle | Transferase(Phosphotransferase); domain 1 |
| 2nd | 1.10.8 | Mainly Alpha | Orthogonal Bundle | Helicase, Ruva Protein; domain 3 |
| 2nd | 1.20.120 | Mainly Alpha | Up-down Bundle | Four Helix Bundle (Hemerythrin (Met), subunit A) |
| 2nd | 1.20.1250 | Mainly Alpha | Up-down Bundle | Growth Hormone; Chain: A; |
| 2nd | 1.20.5 | Mainly Alpha | Up-down Bundle | Single alpha-helices involved in coiled-coils or other helix-helix interfaces |
| 2nd | 1.20.58 | Mainly Alpha | Up-down Bundle | Methane Monooxygenase Hydroxylase; Chain G, domain 1 |
| 2nd | 1.25.10 | Mainly Alpha | Alpha Horseshoe | Leucine-rich Repeat Variant |
| 2nd | 1.25.40 | Mainly Alpha | Alpha Horseshoe | Serine Threonine Protein Phosphatase 5, Tetratricopeptide repeat |
| 2nd | 2.130.10 | Mainly Beta | 7 Propeller | Methylamine Dehydrogenase; Chain H |
| 2nd | 2.30.30 | Mainly Beta | Roll | SH3 type barrels. |
| 2nd | 2.40.50 | Mainly Beta | Beta Barrel | OB fold (Dihydrolipoamide Acetyltransferase, E2P) |
| 2nd | 2.60.120 | Mainly Beta | Sandwich | Jelly Rolls |
| 2nd | 2.60.40 | Mainly Beta | Sandwich | Immunoglobulin-like |
| 2nd | 3.20.20 | Alpha Beta | Alpha-Beta Barrel | TIM Barrel |
| 2nd | 3.30.200 | Alpha Beta | 2-Layer Sandwich | Phosphorylase Kinase; domain 1 |
| 2nd | 3.30.40 | Alpha Beta | 2-Layer Sandwich | Herpes Virus-1 |
| 2nd | 3.30.420 | Alpha Beta | 2-Layer Sandwich | Nucleotidyltransferase; domain 5 |
| 2nd | 3.30.70 | Alpha Beta | 2-Layer Sandwich | Alpha-Beta Plaits |
| 2nd | 3.40.30 | Alpha Beta | 3-Layer(aba) Sandwich | Glutaredoxin |
| 2nd | 3.40.50 | Alpha Beta | 3-Layer(aba) Sandwich | Rossmann fold |
| 2nd | 3.50.50 | Alpha Beta | 3-Layer(bba) Sandwich | FAD/NAD(P)-binding domain |
| 2nd | 3.80.10 | Alpha Beta | Alpha-Beta Horseshoe | Leucine-rich repeat, LRR (right-handed beta-alpha superhelix) |
| intermediate | 1.10.1070 | Mainly Alpha | Orthogonal Bundle | Phosphatidylinositol 3-kinase Catalytic Subunit; Chain A, Domain 5 |
| intermediate | 1.10.132 | Mainly Alpha | Orthogonal Bundle | Topoisomerase I; Chain A, domain 4 |
| intermediate | 1.10.20 | Mainly Alpha | Orthogonal Bundle | Histone, subunit A |
| intermediate | 1.10.220 | Mainly Alpha | Orthogonal Bundle | Annexin V; domain 1 |
| intermediate | 1.10.30 | Mainly Alpha | Orthogonal Bundle | DNA Binding (I), subunit A |
| intermediate | 1.10.730 | Mainly Alpha | Orthogonal Bundle | Isoleucyl-tRNA Synthetase; Domain 1 |
| intermediate | 1.20.1050 | Mainly Alpha | Up-down Bundle | Glutathione S-transferase Yfyf (Class Pi); Chain A, domain 2 |
| intermediate | 1.20.1110 | Mainly Alpha | Up-down Bundle | Calcium-transporting ATPase, transmembrane domain |
| intermediate | 1.20.1740 | Mainly Alpha | Up-down Bundle | Amino acid/polyamine transporter I |
| intermediate | 1.50.10 | Mainly Alpha | Alpha/alpha barrel | Glycosyltransferase |
| intermediate | 2.160.20 | Mainly Beta | 3 Solenoid | Pectate Lyase C-like |
| intermediate | 2.70.150 | Mainly Beta | Distorted Sandwich | Calcium-transporting ATPase, cytoplasmic transduction domain A |
| intermediate | 3.10.110 | Alpha Beta | Roll | Ubiquitin Conjugating Enzyme |
| intermediate | 3.30.1360 | Alpha Beta | 2-Layer Sandwich | Gyrase A; domain 2 |
| intermediate | 3.30.310 | Alpha Beta | 2-Layer Sandwich | TATA-Binding Protein |
| intermediate | 3.30.559 | Alpha Beta | 2-Layer Sandwich | Chloramphenicol Acetyltransferase |
| intermediate | 3.40.1110 | Alpha Beta | 3-Layer(aba) Sandwich | Calcium-transporting ATPase, cytoplasmic domain N |
| intermediate | 3.40.140 | Alpha Beta | 3-Layer(aba) Sandwich | Cytidine Deaminase; domain 2 |
| intermediate | 3.40.250 | Alpha Beta | 3-Layer(aba) Sandwich | Oxidized Rhodanese; domain 1 |
| intermediate | 3.60.15 | Alpha Beta | 4-Layer Sandwich | Metallo-beta-lactamase; Chain A |
| intermediate | 3.90.640 | Alpha Beta | Alpha-Beta Complex | Actin; Chain A, domain 4 |
| intermediate | 3.90.79 | Alpha Beta | Alpha-Beta Complex | Nucleoside Triphosphate Pyrophosphohydrolase |
| 1st | 1.10.1500 | Mainly Alpha | Orthogonal Bundle | Probable Glutaminase Ybgj; Chain: A, domain 2 |
| 1st | 1.10.60 | Mainly Alpha | Orthogonal Bundle | Diphtheria Toxin Repressor; domain 2 |
| 1st | 2.170.8 | Mainly Beta | Beta Complex | Phosphoenolpyruvate Carboxykinase; domain 2 |
| 1st | 2.90.10 | Mainly Beta | Orthogonal Prism | Agglutinin, subunit A |
| 1st | 3.30.10 | Alpha Beta | 2-Layer Sandwich | Trypsin Inhibitor V; Chain A |
| 1st | 3.30.2170 | Alpha Beta | 2-Layer Sandwich | archaeoglobus fulgidus dsm 4304 fold |
| 1st | 3.40.1080 | Alpha Beta | 3-Layer(aba) Sandwich | Glutaconate Coenzyme A-transferase |
| 1st | 3.40.109 | Alpha Beta | 3-Layer(aba) Sandwich | NADH Oxidase |
| 1st | 3.40.449 | Alpha Beta | 3-Layer(aba) Sandwich | Phosphoenolpyruvate Carboxykinase; domain 1 |
| 1st | 3.90.330 | Alpha Beta | Alpha-Beta Complex | Nitrile Hydratase; Chain A |

**Supplementary Table 1. List of CAT ID, Class, Architecture and Topology of the folds in each core category.**

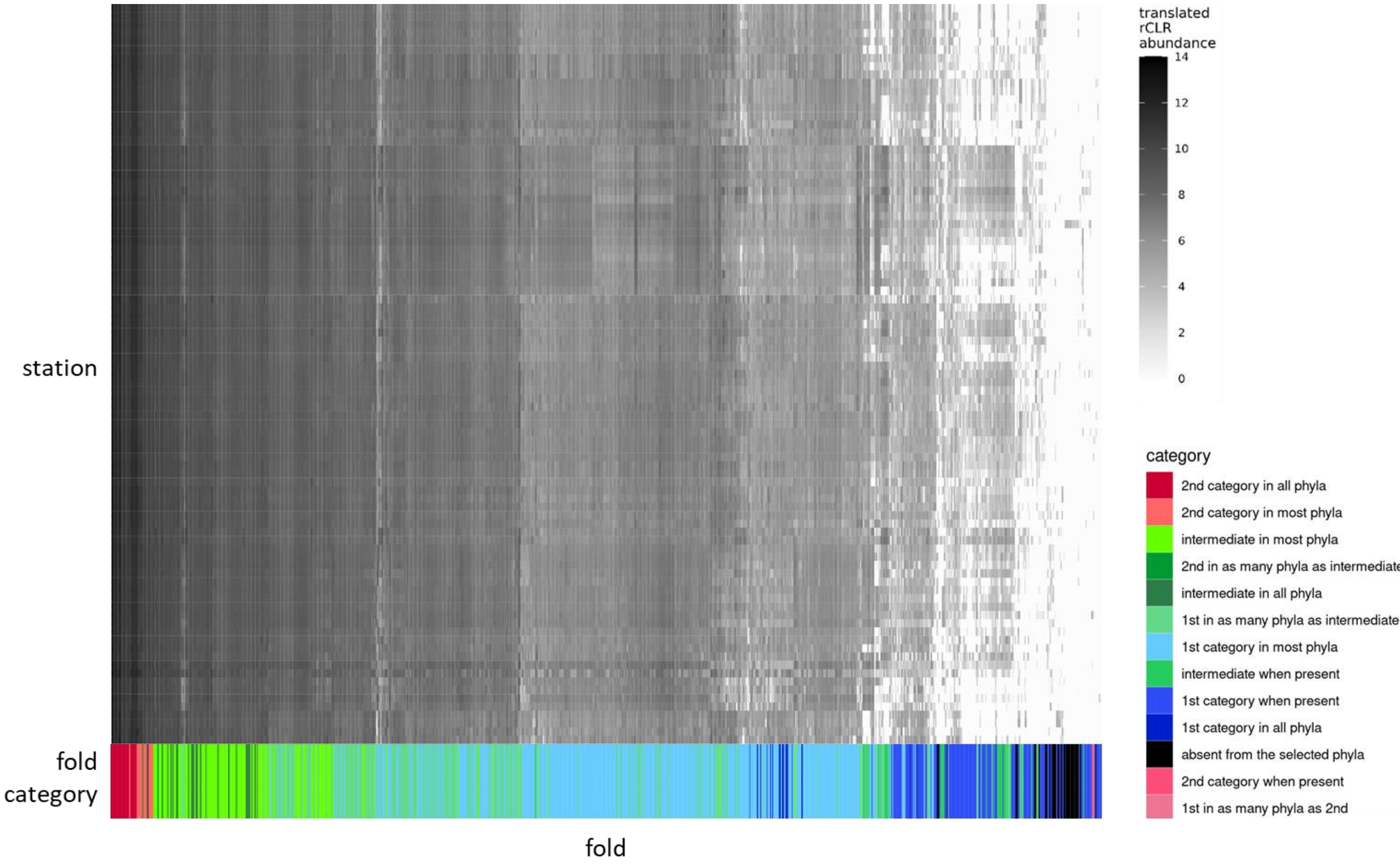


**Supplementary Figure 14. Abundance category of the folds from Fig. 1A.** Fig. 1A with an additional layer on the x-axis indicating the average membership (measured with the six phyla of the previous analyses) of the folds to each abundance category. “in all phyla”: the fold belongs to the category in the six phyla; “in most phyla”: the fold belongs to the category in at least four phyla; “when present”: the fold is not found in at least one of the six phyla.

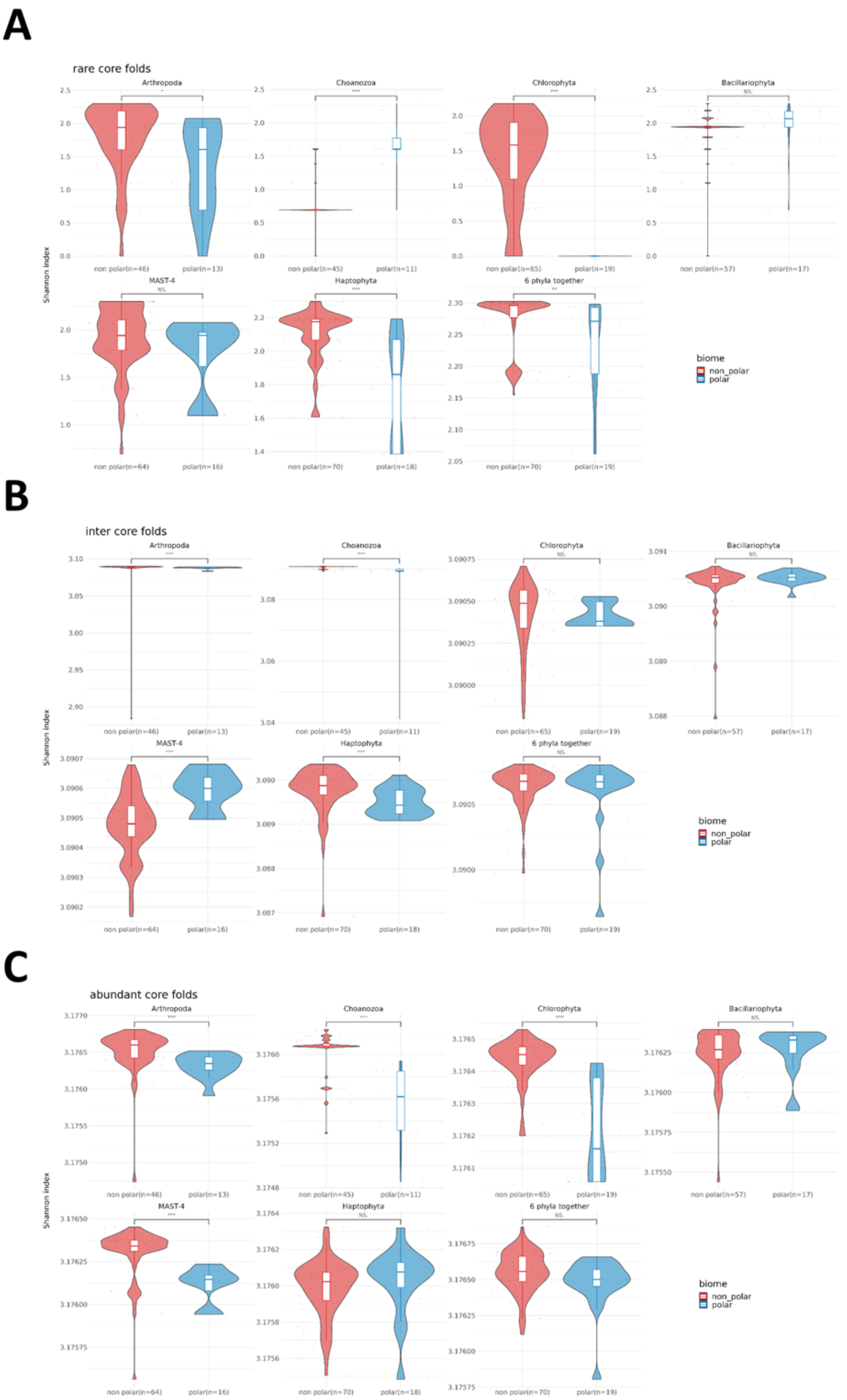


**Supplementary Figure 15. Alpha-diversity differences of fold composition between polar and nonpolar stations per phylum and core of each abundance category.** The α-diversity is measured with the Shannon index. For each plot, from left to right and top to bottom: Arthropoda, Choanoflagellata, Chlorophyta, Bacillariophyta, MAST-4, Haptophyta and the six phyla together. Statistical significance of the differences between values of α-diversity were estimated using a Wilcoxon test (***: $p$-value < 0.01; **: $p$-value < 0.05; *: $p$-value < 0.1; NS: $p$-value ≥ 0.1). TO stations at latitude above 60° North or South are considered as polar (blue violin), the other are nonpolar (red violin). The value in brackets in the x-axis of each facet indicates the number of stations with folds belonging to the phylum in question. **(A)** Folds from the core of the rare category. **(B)** Folds from the core of the intermediate category. **(C)** Folds from the core of the abundant category.

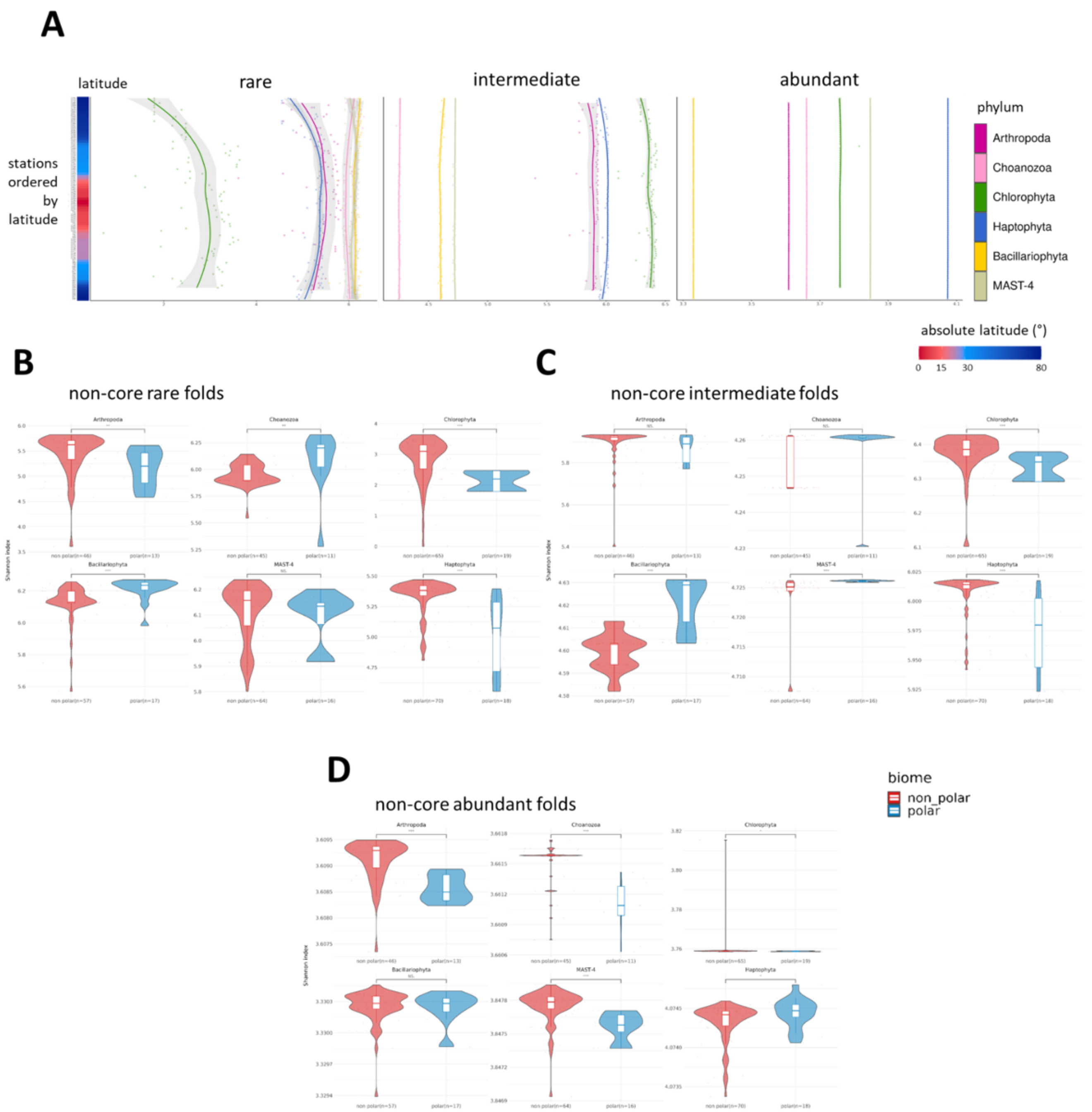


**Supplementary Figure 16. Alpha-diversity of the fold composition of TO stations per abundance category and phylum. (A)** Shannon index in the different fold categories (rare: left; intermediate: centre; abundant: right) per phylum. Stations ordered by latitude are on the y-axis. Shannon index values are indicated on the x-axis. Each colour represents a phylum. The grey zone surrounding the lines indicate the standard error. **(B-C-D)** Differences of Shannon index values between polar and nonpolar stations, measured with the folds of each category. For each plot, from left to right and top to bottom: Arthropoda, Choanoflagellata, Chlorophyta, Bacillariophyta, MAST-4 and Haptophyta. Statistically significant differences between values of α-diversity were estimated using a Wilcoxon test (***: $p$-value < 0.01; **: $p$-value < 0.05; *: $p$-value < 0.1; NS: $p$-value ≥ 0.1). TO stations at latitude above 60°N or S are considered as polar (blue violin), the others are non-polar (red violin). The value in brackets in the x-axis of each facet indicates the number of stations with folds belonging to the phylum in question. **(B)** Folds from the rare category. **(B)** Folds from the intermediate category. **(C)** Folds from the abundant category.

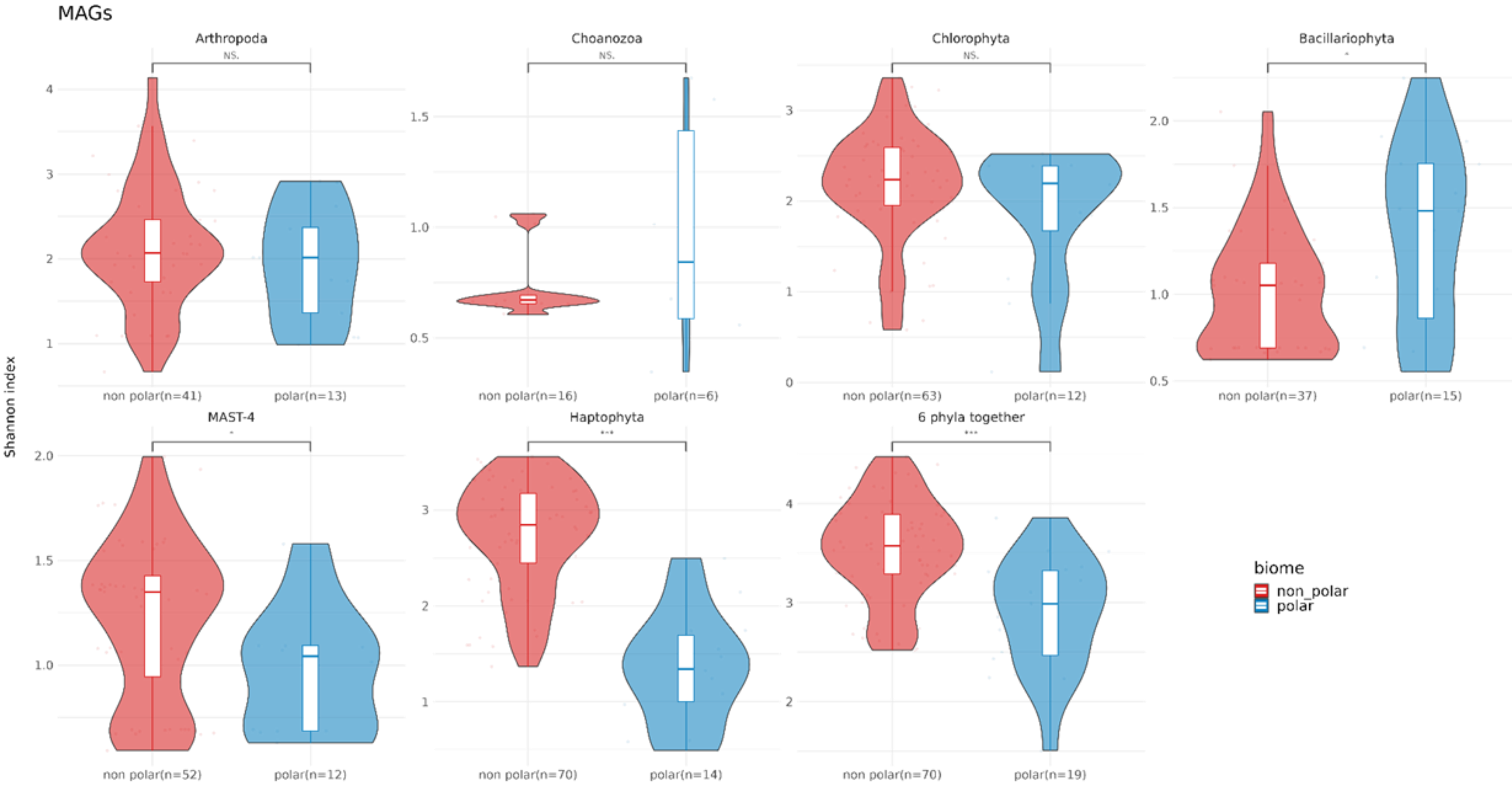


**Supplementary Figure 17. Alpha-diversity differences of MAG composition between polar and non-polar stations per phylum.** The α-diversity is measured with the Shannon index. From left to right and top to bottom: Arthropoda, Choanoflagellata, Chlorophyta, Bacillariophyta, MAST-4, Haptophyta and the 6 phyla together. Statistically significant differences between values of α-diversity were estimated using a Wilcoxon test (***: *p*-value < 0.01; **: *p*-value < 0.05; *: *p*-value < 0.1; NS: *p*-value ≥ 0.1). TO stations at latitude above 60°N or S are considered as polar (blue violin), the others are nonpolar (red violin). The value in brackets in the x-axis of each facet indicates the number of stations with MAGs belonging to the phylum in question.

| category \ phylum | | Arthropoda | Choanozoa | Chlorophyta | Haptophyta | Bacillariophyta | MAST-4 | 6 phyla together |
|---|---|---|---|---|---|---|---|---|
| core | rare | NS | *** | *** | *** | NS | NS | NS |
| | intermediate | *** | *** | NS | *** | NS | *** | NS |
| | abundant | *** | *** | *** | NS | NS | *** | NS |
| non-core | rare | NS | NS | *** | *** | *** | NS | |
| | intermediate | NS | NS | *** | *** | *** | *** | / |
| | abundant | *** | *** | NS | NS | NS | *** | |
| MAGs | | NS | NS | NS | *** | NS | NS | *** |

**Supplementary Table 2. Result of the Wilcoxon tests on the α-diversity values between polar and non-polar stations measured with folds per fold category and phylum or MAGs per phylum.** Summary of the results from Supplementary Fig.14 to 16. Cells in red with “NS” correspond to non-significant Wilcoxon test (*p*-value ≥ 0.01). Cells in green with “***” correspond to significant Wilcoxon test (*p*-value < 0.01).

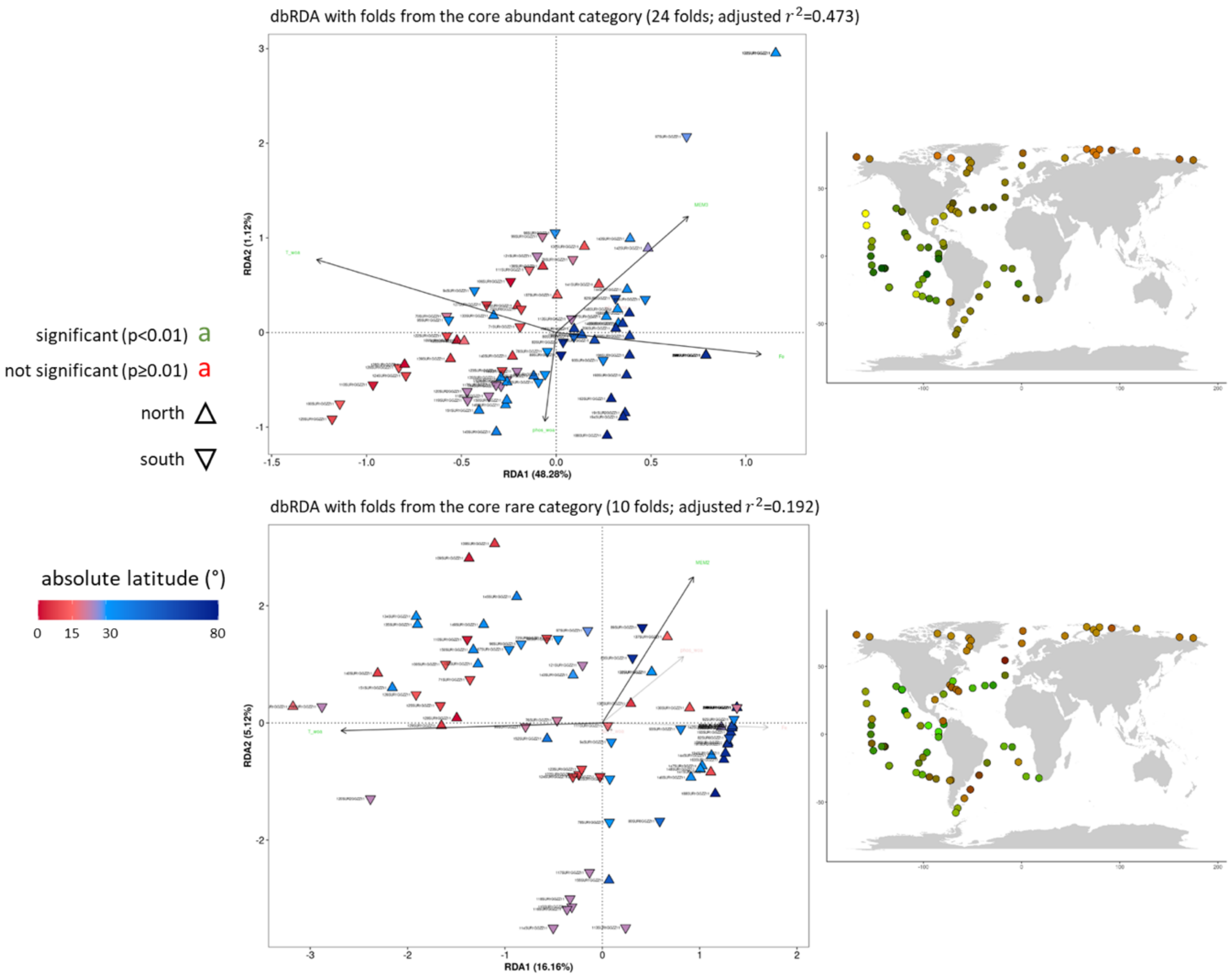


**Supplementary Figure 18. Structuring of the biogeographical distribution of the folds from the cores of the abundant and rare categories in Chlorophyta.** On the top: folds from the core of the abundant category. On the bottom: folds from the core of the rare category. The two distance-based ReDundancy Analyses (dbRDAs) biplots are represented on the left. Each subtitle indicates the fold category and the number of folds in the corresponding core in brackets, followed by the value of the adjusted coefficient of determination $r^2$ (proportion of variance of the fold matrix explained by the selected variables). In the biplots, the coloured triangles represent the stations. Their colour correspond to the absolute latitude. Triangles pointing upwards correspond to stations above the equator. Downward-pointing triangles correspond to stations below the equator. Black arrows correspond to statistically significant explanatory variables, and point toward names in green. They are transparent otherwise and point toward names in red. MEM: Moran Eigenvector Map. T_woa: mean annual SST extracted from the WOA [2]. Sd_T_woa: standard deviation of the mean annual SST extracted from the WOA. Phos_woa: Phosphate concentration extracted from the WOA. Fe: Iron concentration. The coordinates of the stations in the two first dimensions of the dbRDA space were converted to a Red-Green (RG) code that was used to colour the corresponding stations in the world maps.

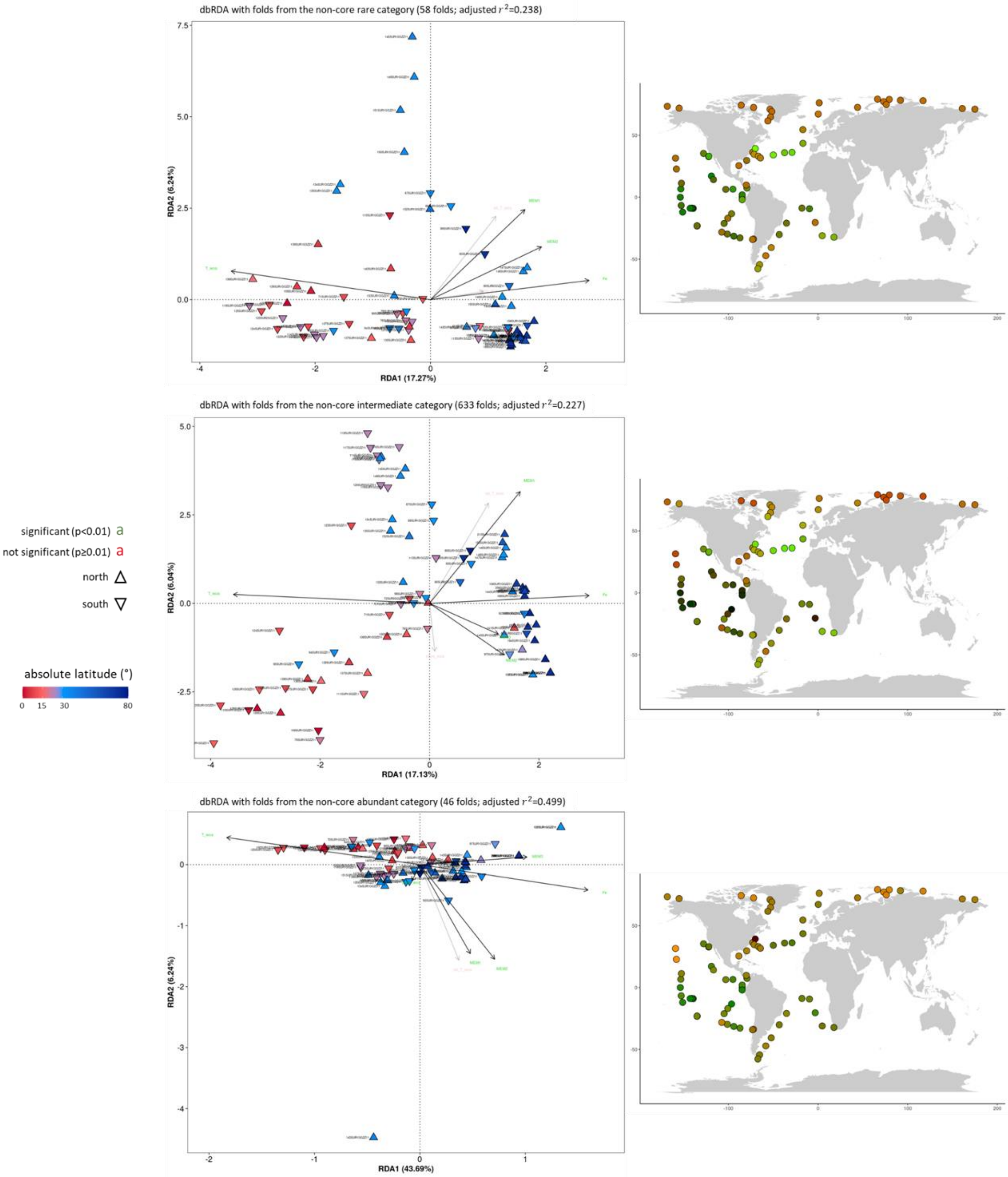


**Supplementary Figure 19. Structuring of the biogeographical distribution of the folds from the three abundance categories in Chlorophyta.** From top to bottom: folds from the rare category, folds from the intermediate category and folds from the abundant category. The three dbRDA biplots are on the left. Each subtitle indicates the fold category and the number of folds in that category in brackets, followed by the value of the $r^2$. In the biplots, the coloured triangles represent the stations. Their colour correspond to the absolute latitude. Triangles pointing upwards correspond to stations above the equator. Downward-pointing triangles correspond to stations below the equator. Black arrows correspond to statistically significant explanatory variables, and point toward names in green. They are transparent otherwise and point toward names in red. The coordinates of the stations in the two first dimensions of the dbRDA space were converted to a RG code that was used to colour the corresponding stations in the world maps.

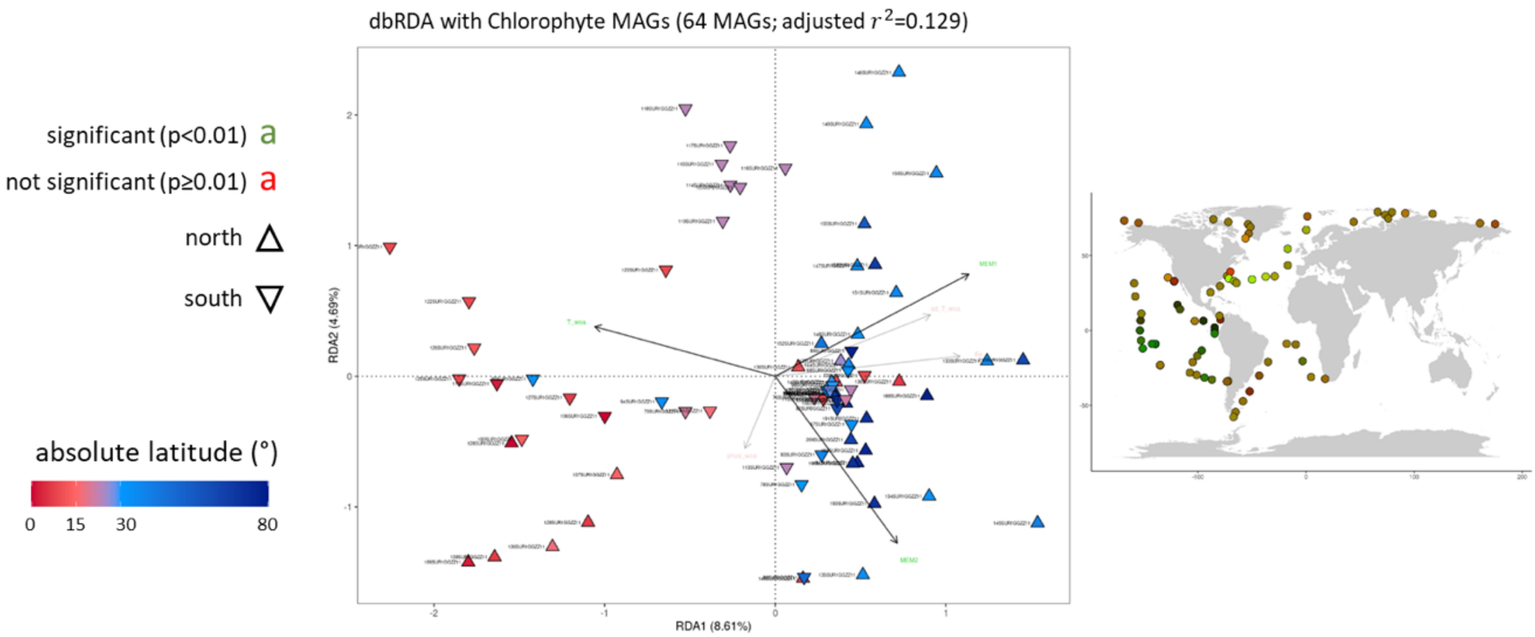


**Supplementary Figure 20. Structuring of the biogeographical distribution of the Chlorophyte MAGs.** In the subtitle, the number of Chlorophyte MAGs is indicated in brackets, followed by the $r^2$. In the biplot, the coloured triangles represent the stations. Their colour correspond to the absolute latitude. Triangles pointing upwards correspond to stations above the equator. Downward-pointing triangles correspond to stations below the equator. Black arrows correspond to statistically significant explanatory variables, and point toward names in green. They are transparent otherwise and point toward names in red. The coordinates of the stations in the two first dimensions of the dbRDA space were converted to a RG code that was used to colour the corresponding stations in a world map.

| category | phylum | Arthropoda | Choanozoa | Chlorophyta | Haptophyta | Bacillariophyta | MAST-4 |
|---|---|---|---|---|---|---|---|
| core | rare | 14,89 | 22,6 | 16,16 | 12,69 | 17,29 | 10,68 |
| | intermediate | 15,1 | 29,17 | 44,8 | 34,46 | 10,15 | 12,82 |
| | abundant | 9,1 | 47,16 | 48,28 | 25,77 | 13,89 | 7,32 |
| non-core | rare | 10,35 | 23,72 | 17,27 | 11,3 | 10,3 | 12,49 |
| | intermediate | 12,25 | 42,18 | 17,13 | 18,28 | 14,51 | 33,32 |
| | abundant | 9,32 | 44,63 | 43,69 | 22,49 | 13,74 | 9,49 |
| MAGs | | 8,38 | 0 | 8,61 | 8,4 | 6,39 | 10,59 |

| category | phylum | Arthropoda | Choanozoa | Chlorophyta | Haptophyta | Bacillariophyta | MAST-4 |
|---|---|---|---|---|---|---|---|
| core | rare | 2,41 | 3,06 | 5,12 | 2,02 | 3,5 | 5,01 |
| | intermediate | 2,66 | 8,01 | 6,12 | 4,02 | 2,8 | 3,08 |
| | abundant | 4,42 | 1,95 | 1,12 | 6,71 | 2,21 | 3,15 |
| non-core | rare | 2,29 | 6,1 | 6,24 | 3,28 | 4,12 | 5,81 |
| | intermediate | 4,08 | 4,76 | 6,04 | 6,27 | 3,77 | 2,83 |
| | abundant | 3,51 | 2,04 | 6,24 | 4,89 | 2,15 | 3,53 |
| MAGs | | 3,75 | 0 | 4,69 | 6,04 | 4,76 | 6,84 |

**Supplementary Table 3. Percentage of variance explained by the two first axis of the dbRDAs for each phylum, fold category and MAG.** The values are in percentage. Top: first dimension. Bottom: second dimension.

| category \ phylum | | Arthropoda | Choanozoa | Chlorophyta | Haptophyta | Bacillariophyta | MAST-4 |
|---|---|---|---|---|---|---|---|
| **core** | **rare** | 0,132 | 0,215 | 0,192 | 0,113 | 0,196 | 0,137 |
| | **intermediate** | 0,139 | 0,337 | 0,491 | 0,372 | 0,106 | 0,14 |
| | **abundant** | 0,076 | 0,456 | 0,473 | 0,303 | 0,126 | 0,066 |
| **non-core** | **rare** | 0,094 | 0,288 | 0,238 | 0,132 | 0,142 | 0,191 |
| | **intermediate** | 0,141 | 0,445 | 0,227 | 0,23 | 0,171 | 0,356 |
| | **abundant** | 0,073 | 0,431 | 0,499 | 0,251 | 0,125 | 0,095 |
| **MAGs** | | 0,097 | / | 0,129 | 0,169 | 0,088 | 0,175 |

**Supplementary Table 4. Determination coefficient of the dbRDAs for each phylum, fold category and MAGs.**

| category \ phylum | | Arthropoda | Choanozoa | Chlorophyta | Haptophyta | Bacillariophyta | MAST-4 |
|---|---|---|---|---|---|---|---|
| **core** | **rare** | T, Fe | T, P | MEM2, T | T | T, Fe, P | MEM1, T |
| | **intermediate** | T | T, Fe, P | MEM1, MEM3, T, Fe | T, sd(T), P | T, sd(T) | T, P |
| | **abundant** | T | T, Fe, P | MEM3, T, Fe, P | T, sd(T), P | T | / |
| **non-core** | **rare** | T, Fe | T, sd(T), P | MEM1, MEM2, T, Fe | T, sd(T), Fe, P | MEM1, T, Fe, P | MEM1, T, Fe, P |
| | **intermediate** | T, sd(T), Fe | T, Fe, P | MEM1, MEM2, MEM3, T, Fe | T, P | T, Fe, P | T, P |
| | **abundant** | T | T, Fe, P | MEM1, MEM2, MEM3, T, Fe, P | T, sd(T), P | T | T, P |
| **MAGs** | | MEM1, T | / | MEM1, MEM2, T | MEM1, MEM2, MEM3, T, P | T, sd(T), P | MEM1, T, P |

**Supplementary Table 5. Statistically significant explanatory variables for each phylum, fold category and MAGs.** In yellow: MEM. In red: T (mean annual SST from WOA [2]). In purple: sd(T) (standard deviation of the mean annual SST from WOA). In grey: Fe. In green: P (phosphate concentration from WOA). No statistically significant variable was found for the core abundant folds in MAST-4 and there are too few Choanoflagellate MAGs to perform a dbRDA.
